# Hemodynamic-Structural Multi-Fidelity Assessment of Endovascular Flow Diversion Strategies for Cerebral Aneurysm Treatment: An Integrated Computational and Experimental Framework

**DOI:** 10.64898/2026.10.13.758431

**Authors:** Dipu Roy, Farah Sadia Antara, Shama Abdullah Moshrur, Maruf Hasan, Kaiissar Mannoor, Muhammad Shattique, Jafrin Sultana

**Affiliations:** Department of Biomedical Engineering, Military Institute of Science and Technology, Dhaka 1216, Bangladesh

**Keywords:** Cerebral aneurysm, Computational Fluid Dynamics, Wall Shear Stress, Flow Diversion, Finite Element Analysis, Additive Manufacturing, Hemodynamic

## Abstract

Subarachnoid hemorrhage after cerebral aneurysm rupture has a 50% first-month fatality rate. Endovascular flow diversion is the primary treatment modality. However, no prior study has compared stents, coils, and the Woven Endo-Bridge hemodynamically integrating hemodynamic results with structural device behavior in one framework. An aneurysm model was reconstructed from computed tomography imaging of a patient, and computational fluid dynamics evaluated multiple hemodynamic metrics across three flow-diversion strategies. Finite element analysis assessed stent geometries with 50%, 55%, and 60% porosity in 316L stainless steel, poly-L-lactic acid, and nitinol. Universal testing machine tests validated the models with printed polylactic acid and thermoplastic polyurethane analogues. Dimensional and elastic-tube analyses tested scale equivalence between the FEA and UTM geometries and the rigid-wall CFD assumption. Patient-specific CFD confirmed a high-risk hemodynamic profile. The stent yielded the lowest peak intra-sac velocity, greatest pressure reduction, near-zero wall shear stress, the only reduction in time-averaged wall shear stress, cycle-averaged TAWSS, and the largest peak-systolic reductions in velocity, WSS, and vorticity. OSI and RRT rose for all three treated configurations, consistent with the hemodynamic signature of flow diversion. FEA reproduced the expected stiffness hierarchy (Steel > Nitinol > PLLA), and TPU was the closest surrogate for Nitinol’s elastic recovery. Scaled specimens matched native-scale stiffness within 2.3%. Predicted radial distension was orders of magnitude below native wall values, supporting rigid-wall CFD. Stent-assisted diversion demonstrated the most favorable hemodynamic profile. A 50%-porosity Nitinol stent best matched target elastic mechanics, while 316L stainless steel provided the greatest resistance to radial compression.

## 1. Introduction

Around 3.2% of people have a cerebral aneurysm [1], and every year, in the US alone, ruptures cause about 30,000 cases of aneurysmal subarachnoid hemorrhage (aSAH) [2]. Within the first month, nearly half the people who suffer an aSAH die [3]. That statistic is the reason why neurovascular teams put so much effort into catching high-risk lesions before they rupture.

Currently, three endovascular approaches dominate the most. Flow-diverting stents sit across the aneurysm neck and reroute blood away from the sac. Coils pack the sac directly and trigger clotting. Intrasaccular devices like the Woven Endo-Bridge (WEB) cuts down flow into the neck without taking up space in the parent artery itself.

Mechanical performance of devices after deployment is solely depended on material choice. The default metal in stent manufacturing is 316L stainless steel due to its corrosion resistance property and low cost, though bare 316L surfaces still carry a reported 15-20% restenosis risk, which has pushed recent work toward nanostructured surface treatments improving endothelization and decreases platelet adhesion [4]. Nitinol has replaced stainless steel in self-expanding neurovascular platforms. Stress-induced martensitic transformation of Nitinol gives it superelastic recovery of up to 8% strain, letting a device compress into a delivery catheter and spring back to its original shape [5]. Bioresorbable poly-L-lactic acid (PLLA) scaffolds are also under active development. PLLA degrades fully over time, but its lower stiffness relative to metals and its slow, acid-producing degradation limit strut thickness along with long-term mechanical support [6]. Polylactic acid (PLA) is a related but distinct polymer, which is increasingly used as a stent coating and a fused-deposition-modeling surrogate, since its stiffness approximates PLLA’s mechanical response at a much lower manufacturing cost [7]. Thermoplastic polyurethane (TPU) has become a common material for 3D-printed vascular phantoms and stent-graft prototypes. Its biphasic hard-soft segment structure gives it a gradual, elastically recoverable compression response that mirrors Nitinol’s qualitative behavior, although the two materials differ by orders of magnitude in elastic modulus [8]. These five materials span the practical design space for current and emerging flow-diversion devices from rigid metals to compliant polymers. This range motivates the structural comparison undertaken in the present study.

For characterizing aneurysm hemodynamics, computational fluid dynamics (CFD) has become the go-to method, and it doubles as a way to test how much a given device actually changes the flow. Researchers lean on wall shear stress (WSS), time-averaged WSS (TAWSS), relative residence time (RRT), and oscillatory shear index (OSI) as proxies for how likely an aneurysm is to grow or rupture [9,10]. The pattern that keeps showing up. High WSS at the neck seems to kick off the endothelial damage that starts an aneurysm forming, while low WSS paired with high OSI and RRT keeps the inflammation going that drives growth toward rupture [11]. Although, CFD-based device assessment runs into two real limitations in clinical practice. First, virtually all CFD aneurysm studies use rigid-wall models. These models demonstrably overestimate WSS relative to compliant-wall fluid– structure interaction (FSI) models [12]. Second, studies almost never characterize implanted device structural behavior alongside hemodynamic performance. Clinically documented failure modes include stent fracture [13] and coil compaction [14].

A third gap concerns geographic representation. Published CFD aneurysm cohorts contain few South Asian patients [15]. This gap persists despite documented differences in aneurysm morphology and hypertension burden between Asian and Western populations [16,17].

The present study addresses the three identified deficits through a multi-phase integrated framework. We reconstructed a patient-specific aneurysm from imaging data (BME-REC-2026-09-01) obtained at the National Institute of Neurosciences and Hospital, Dhaka, Bangladesh. We performed computational fluid dynamics (CFD) to quantify hemodynamic risk. We then compared stent, coil, and WEB strategies in an idealized aneurysm model under identical pulsatile conditions using seven hemodynamic parameters. The patient-specific reconstruction confirmed a high-risk flow regime (TAWSS = 6.68 Pa, OSI = 0.180,

RRT = 2.63 Pa ¹ in Section 3.1). This result directly established the study’s clinical motivation. We used an idealized geometry for device comparison. Parent vessel tortuosity, aneurysm neck taper, and other case-specific features vary independently of device design. Retaining these features in a four-way comparison would confound attribution of hemodynamic differences to device configuration rather than anatomy. The idealized model isolated device-intrinsic mechanics from patient-specific anatomical variability. The patient-specific reconstruction alone cannot provide this isolation. We then performed static structural finite element analysis (FEA) on three custom stent geometries using three clinically representative material models. We established the dimensional analysis conditions for mechanical equivalence between scaled universal testing machine (UTM) specimens and native-scale devices. We applied classical elastic-tube theory to bound the contribution of device radial compliance to intra-aneurysmal flow. This analysis provided a first-order check of the rigid-wall assumption underlying the hemodynamic analysis. We demonstrated the first integrated framework that combines hemodynamic assessment of all three principal device classes with structural characterization of representative prototypes using South Asian patient data.

## 2. Materials and methods

### 2.1. Computational Fluid Dynamics

#### 2.1.1. Patient-Specific Image Acquisition and Geometry Reconstruction

We acquired CT data from a 67-year-old patient (BME-REC-2026-09-01) with an internal carotid artery aneurysm. We received institutional approval from the National Institute of Neurosciences and Hospital, Dhaka, Bangladesh. We acquired approximately 1,050 DICOM-format image slices with a spatial resolution of 512×512 pixels using a SOMATOM Definition AS+ system. We reconstructed the artery using Materialise Mimics. We excluded minor arterial branches. We applied surface smoothing to remove geometric irregularities and exported the finalized geometry in IGES format for import into ANSYS.

#### 2.1.2. Patient-Specific Aneurysmal Artery Simulation

We imported the reconstructed geometry into the computational domain. We simulated physiological flow conditions. We then quantified baseline intra-aneurysmal flow before implementing an endovascular treatment strategy.

#### 2.1.3. Mesh Generation and Mesh Independence Study

A mesh-independence study was performed, and a mesh of 3.7 million elements was selected because further refinement changed WSS by <1% (Supplementary Table 1, Supplementary Figure 1a).

#### 2.1.4. Governing Equations and Blood Rheology

We modeled blood flow with the unsteady, incompressible Navier-Stokes and continuity equations (1) and (2).

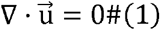

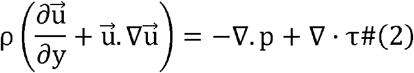

Here, *u* is the velocity vector, *p* is the static pressure, *ρ* is the fluid density, and *τ* is the viscous stress tensor. We modeled blood as an incompressible, laminar, non-Newtonian fluid with density 1,060 kg/m³ and dynamic viscosity 0.0035 Pa·s [18]. We selected the Carreau-Yasuda viscosity model to capture shear-thinning behavior with parameters:

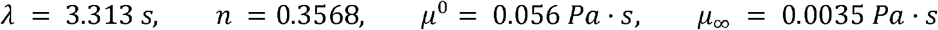

We treated the vessel and device walls as rigid. We imposed a no-slip condition at all luminal surfaces, following standard rigid-wall hemodynamic practice [19]. Section 2.5 derives a closed-form bound for this assumption. The bound demonstrates that radial distension across all nine device configurations is negligible relative to native wall compliance under physiological pulse pressure. We used a pressure-based solver with the SIMPLE algorithm to couple pressure and velocity for the simulations themselves. Pressure and momentum equations were both discretized with second-order upwind schemes, and we required residuals to converge below 10 at every time step.

#### 2.1.5. Boundary Conditions

We prescribed identical and time-dependent boundary conditions for four geometries. At the proximal (inlet) face, we imposed a pulsatile velocity-inlet condition, *v* = *v*(*t*), over one cardiac cycle and a corresponding pulsatile static-pressure condition at the distal (outlet) face (Figure 1). We digitized both waveforms over a 0.8 s cardiac cycle. This cycle corresponds to a resting heart rate of approximately 75 beats per minute and matches internal carotid inflow waveforms from idealized sidewall aneurysm computational fluid dynamics (CFD) studies [20].

**Figure 1:**
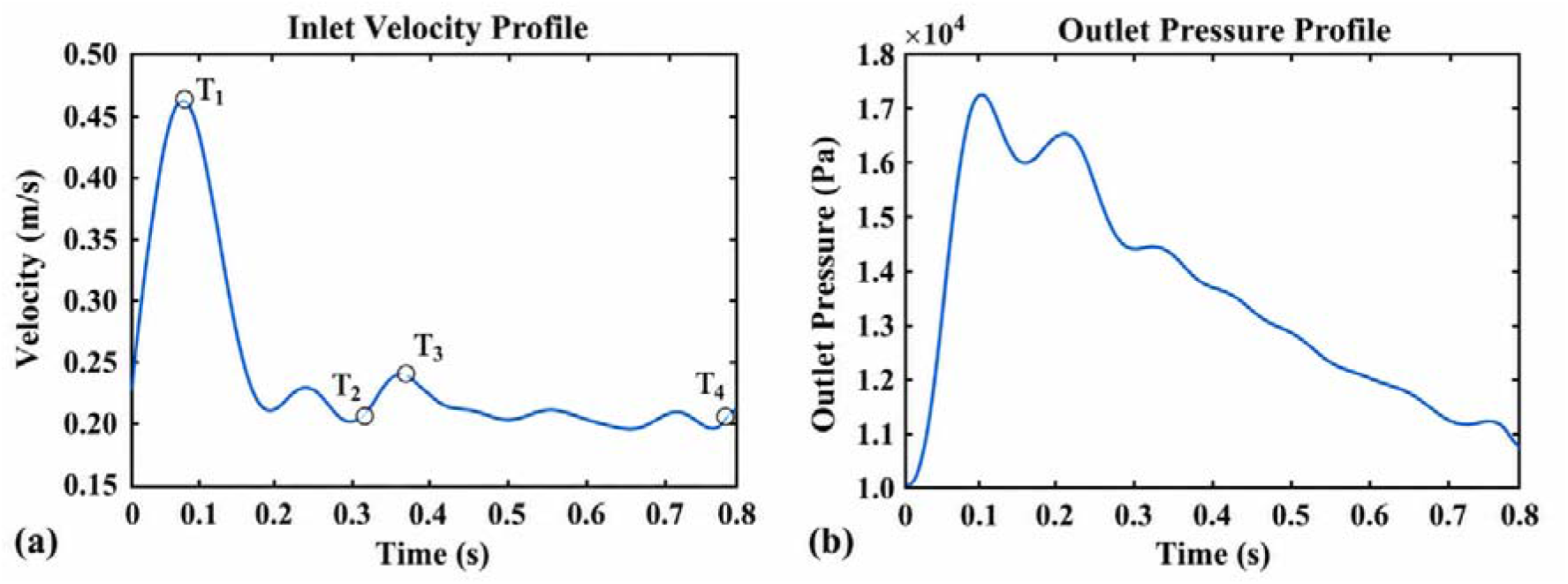
Pulsatile boundary conditions applied identically to all four geometries: (a) inlet velocity profile with T –T marking the characteristic points of the waveform (T = peak systole, T ≈ end diastole), (b) outlet pressure profile over the same 0.8 s cardiac cycle.

The inlet velocity waveform (Figure 1a) contained four characteristic points. T marked the systolic peak: 0.47 m/s at *t* ≈ 0.07 s. T marked a local minimum: approximately 0.21 m/s at *t* ≈ 0.30 s. T marked a secondary maximum: approximately 0.24 m/s at *t* ≈ 0.37 s. T marked a late-diastolic plateau: approximately 0.21 m/s at *t* ≈ 0.75–0.78 s.

The outlet pressure waveform (Figure 1b) rose from a diastolic baseline of ≈ 1.0 × 10 Pa to a systolic peak of ≈ 1.73 × 10 Pa. The pressure peak followed the velocity peak, exhibited a secondary pressure peak of ≈ 1.63 × 10 Pa at *t* ≈ 0.25 s, and decayed gradually to ≈ 1.1 × 10 Pa by the end of the cycle. We applied a no-slip condition to all rigid walls, including the surfaces of the deployed stent, coil, and WEB devices. We advanced each simulation through 100-time steps of Δt = 0.01 s, spanning approximately 1.25 cardiac cycles from an impulsive initial condition; the digitized waveform (0.8 s) is repeated for the final 0.2 s of the run to sustain the pulsatile boundary condition. Peak systole and end diastole are reported from this single computed cycle. Cycle-to-cycle periodicity was not separately verified, a limitation addressed in Section 3.10.

#### 2.1.6. Hemodynamic Parameters

We evaluated seven hemodynamic parameters for each computational fluid dynamics (CFD) configuration. Wall shear stress (WSS) is the tangential force per unit area exerted by blood on the vessel wall. We computed the time-averaged wall shear stress (TAWSS) by equation (3).

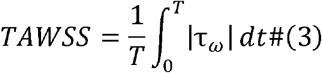

TAWSS gives the cycle-average magnitude of the instantaneous WSS vector over the cardiac period *T*. The oscillatory shear index is calculated by the equation (4).

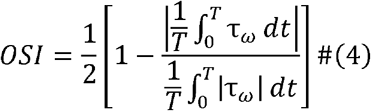

quantifies WSS directional oscillation over the cardiac cycle. OSI ranges from 0 to 0.5, where 0 and 0.5 denotes to fully unidirectional shear and purely oscillatory, net-zero directional shear respectively. The relative residence time is calculated by the equation (5).

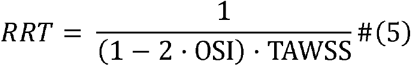

combines TAWSS and OSI into a relative index of near-wall blood residence time. RRT rises sharply under three conditions: low TAWSS, OSI near 0.5, and the combination of these. RRT therefore detects near-stagnant, highly disturbed near-wall flow [21].

RRT is computed pointwise from local TAWSS and OSI and then area-averaged across the wall, and that this is therefore not equivalent to evaluating the formula at the area-averaged TAWSS and OSI values.

#### 2.1.7. Idealized Cerebral Aneurysm-Device Models

We developed an idealized side-wall cerebral aneurysm model. We evaluated four endovascular treatment strategies under controlled geometric and physiological conditions. This idealized geometry, rather than the patient-specific reconstruction in Section 2.1.1, supports the device-to-device comparison that follows. Parent-vessel tortuosity, aneurysm neck taper, and other anatomical features specific to the single reconstructed patient case vary independently of device design. Retaining these features would confound attribution of hemodynamic differences to device configuration. The comparison therefore isolates device effects from anatomical effects. The idealized model separates device-intrinsic mechanics from this patient-specific anatomical variability. The patient-specific reconstruction (Section 3.1) separately demonstrates that the underlying flow regime motivating treatment is clinically high-risk. The computational model used a straight cylindrical parent artery with an internal diameter of 4 mm, representative of the intracranial internal carotid artery, which is one of the most common locations for side-wall cerebral aneurysms [22,23]. A saccular aneurysm was constructed on the lateral wall of the parent vessel. Table 1 defines a medium-sized, wide-neck intracranial aneurysm, consistent with the morphological characteristics reported in clinical and computational studies [22–24].

**Table 1.** The detailed geometric specifications of the stent, coil and WEB.

| Parameter | Dimension |
| --- | --- |
| Parent Vessel Diameter | 4 mm |
| Aneurysm Neck Diameter | Elliptical, 2 mm × 4.5 mm |
| Aneurysm Dome Height | 7.6 mm |
| Aneurysm Dome Diameter | 8 mm |
| Coil Wire Diameter | 0.1 mm |
| Coil and WEB Diameter | 8 mm |
| Stent Strut Thickness | 0.08 mm |

We developed four computational models: (i) an untreated aneurysm, (ii) a stent-treated aneurysm, (iii) a coil-treated aneurysm, and (iv) a Woven Endo-Bridge (WEB)-treated aneurysm. The parent vessel geometry, aneurysm dimensions, neck morphology, and computational domain were kept identical in all four models to ensure that the implanted device is the only variable influencing the hemodynamic response.

We implemented the stent-treated model (Figure 2b) with a braided flow-diverting stent across the aneurysm neck. We aligned the stent with the parent artery’s inner surface. The stent geometry approximated commercially available intracranial flow-diverting devices [25].

**Figure 2.**
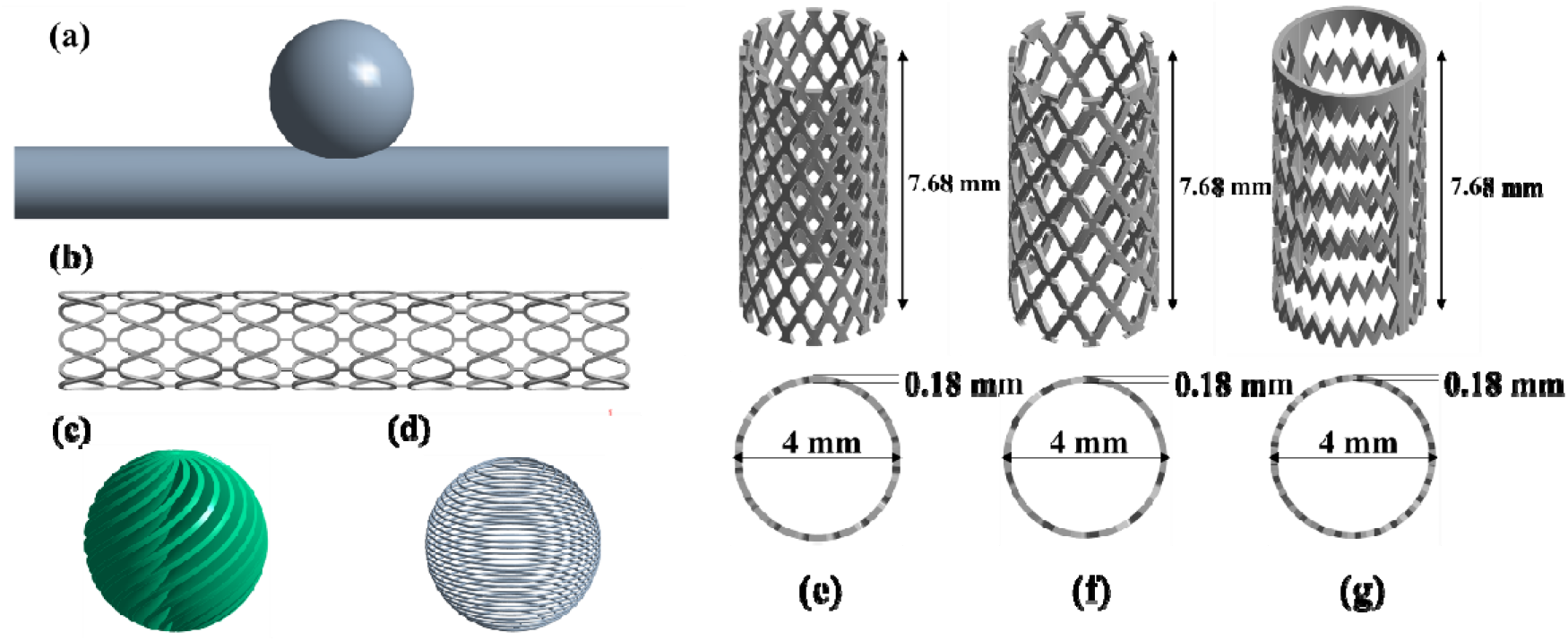
We investigated four computational models: (a) idealized untreated aneurysm; (b) stent; (c) coil; and (d) WEB. We evaluated stent geometries at three porosities: (e) 50%, (f) 55%, and (g) 60%.

We positioned a helical embolization coil (Figure 2c) within the aneurysm sac to simulate endovascular coil embolization. The coil occupied the aneurysm lumen without protruding into the parent artery and disrupted intra-aneurysmal circulation [26].

We implemented the WEB-treated configuration (Figure 2d) with an intrasaccular flow-disruption device at the aneurysm neck. The device remained entirely within the aneurysm sac and disrupted the inflow jet entering the aneurysm, promoting blood flow stagnation and subsequent thrombus formation [27].

We designed all three-dimensional computational models using SolidWorks 2020. After assembling the individual components, we exported the completed geometries to ANSYS Workbench for mesh generation and CFD simulations. We created the parent artery, aneurysm sac, and treatment devices as individual solid bodies before assembling each computational model. We maintained identical geometric dimensions, device-positioning methods, and computational domains across all cases. We therefore compared hemodynamic performance across treatment strategies under controlled geometry and domain conditions.

### 2.2. Finite Element Analysis

#### 2.2.1. Stent Geometry Construction

We developed three stent geometries with target porosities of 50%, 55%, and 60% (Figure 2e–g) using computational modeling methodologies consistent with [28]: P50, P55, and P60 (50%, 55%, and 60% porosity, respectively). All three shared identical outer dimensions: length 7.68 mm, diameter 4 mm, and strut thickness 0.18 mm. Porosity varied with strut spacing and pattern density.

We calculated stent porosity from the strut-structure volume and an equivalent solid-cylinder volume. We defined the fully solid reference as a cylinder with length 7.68 mm and wall thickness 0.18 mm. We calculated porosity using the equation (6).

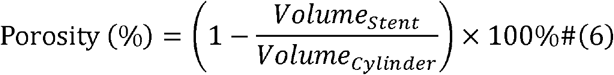

where *Volume_Stent_* denotes the volume of the actual strut geometry, and *Volume_Cylinder_* denotes the volume of the solid reference cylinder. These strut volumes produced porosities of 50.2%, 55.9%, and 59.3%, respectively, within 1% of the target values.

#### 2.2.2. Material Selection

We evaluated three clinical stent materials: 316L stainless steel, poly-L-lactic acid (PLLA), and Nitinol. We modeled 316L stainless steel as linear elastic and isotropic. We assigned Young’s modulus, Poisson’s ratio, and density to each geometry. We modeled PLLA as linear elastic. Static loads remain within its elastic range. PLLA exhibits viscoelasticity under long-term loading [29]. PLLA has lower stiffness than metals [30]. It therefore exhibits greater deformation under identical loads. Nitinol is clinically valued for its super-elastic behavior, driven by a stress-induced phase transformation between austenite and martensite [31]. In this study, however, Nitinol is modeled as linear elastic and isotropic, consistent with 316L stainless steel and PLLA (Supplementary Table 2); this simplification is discussed in Section 3.10.

#### 2.2.3. Mesh Generation and Mesh Independence Study

FEA mesh independence was established based on reaction force, with the selected mesh producing <0.5% deviation from the finest mesh (Supplementary Table 3, Supplementary Figure 1b).

#### 2.2.4. Boundary Conditions and Simulation Setup

We configured the parallel-plate compression test with two rigid plates positioned on either side of the stent shown in Figure 3. The bottom plate was fixed with all degrees of freedom constrained. We prescribed a 1.6 mm displacement for the top plate normal to the stent axis and constrained all other degrees of freedom. Frictional contact was defined between the stent and both plates, with a friction coefficient of 0.2.

**Figure 3.**
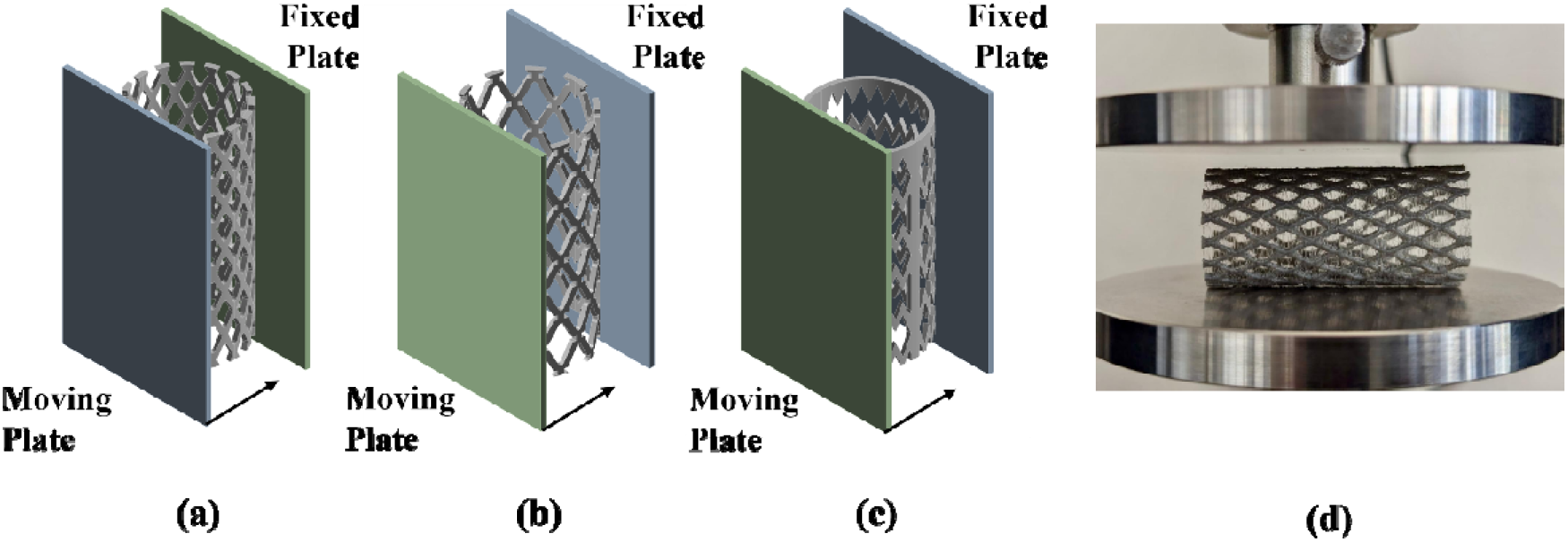
Parallel-plate compression setup for the stent models at (a) 50%, (b) 55%, and (c) 60% porosity, showing the fixed plate, moving plate and displacement direction; (d) Compression test using Universal Testing Machine.

We solved the analysis as a static structural problem with large-deflection kinematics to model geometric nonlinearity in the stent struts under compression. We used one load step with 50 initial sub steps, a minimum of 50, and a maximum of 1000.

#### 2.2.5. Calculated Variables

We extracted four outputs: von Mises equivalent stress, equivalent strain, reaction force, and probe displacement. We recorded equivalent stress and equivalent strain as the maximum values in the stent geometry at full compression. We calculated radial stiffness as the slope of the reaction force versus displacement curve within its initial linear region. For 316L SS, we evaluated this region from 0 to 0.1 mm before the plastic plateau. For Nitinol and PLLA, the response remained linear across the full displacement range. We calculated the slope over 0 to 1.6 mm, defined radial strength as the reaction force at the maximum prescribed displacement, and applied this definition across all nine cases. We did not simulate an unloading cycle.

### 2.3. Experimental Compression Testing

Using a universal testing machine (UTM), we performed parallel-plate compression tests. The machine applied controlled displacement between two rigid plates and recorded the resulting force, which we used to quantify the stent’s radial compression resistance.

#### 2.3.1. Fabrication of Stent Specimens

Our 3D printer cannot fabricate the stent designs used in the finite element analysis (FEA) at their original 4-mm diameter because of its resolution simply isn’t fine enough for physical testing at that scale. Therefore, we scaled all six designs to a length of 48 mm, a diameter of 25 mm, and a strut thickness of 1 mm, while keeping each design’s original porosity (50%, 55%, and 60%). Figure 4 shows the fabricated specimens for all six designs (n = 6). This scaling was necessary for manufacturability and does not represent a proposed clinical dimension.

**Figure 4.**
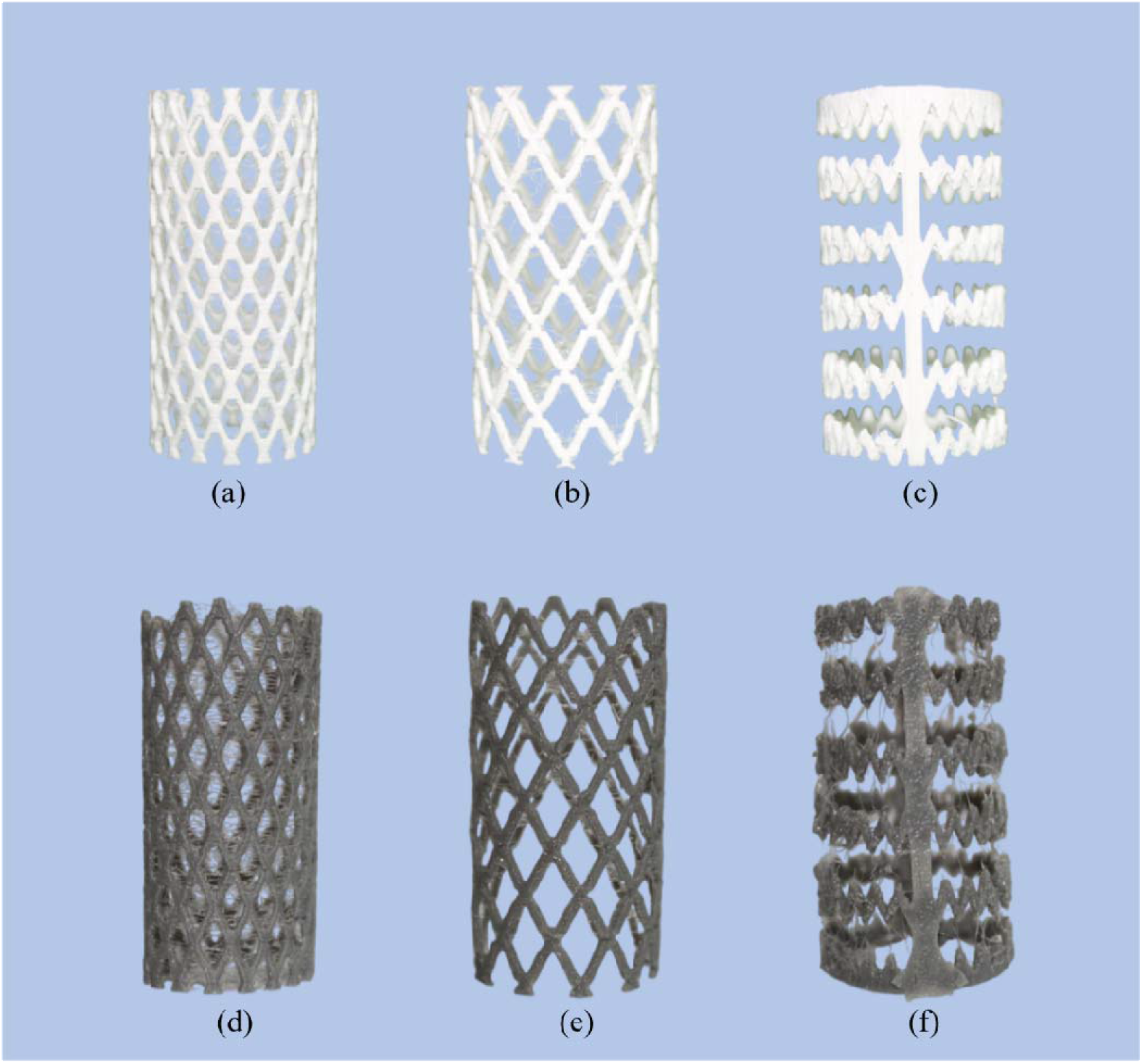
FDM-fabricated stent specimens with different porosity levels using PLA and TPU materials: (a) PLA with 50% porosity, (b) PLA with 55% porosity, (c) PLA with 60% porosity, (d) TPU with 50% porosity, (e) TPU with 55% porosity, and (f) TPU with 60%

Using the available fused deposition modeling (FDM) process, we fabricated the scaled polylactic acid (PLA) and thermoplastic polyurethane (TPU) specimens. FDM parts are inherently anisotropic, with weaker adhesion across layer interfaces than along the print direction [32]. Different print parameters like infill percentage, raster orientation, and nozzle diameter (0.4 mm) directly affect strut strength and stiffness [33]. The printed specimens do not represent the fully dense, isotropic material assumed in the FEA. This difference limits comparisons between experimental and simulated results. Section 3.8 quantifies this effect. Supplementary FEA at the as-printed strut aspect ratio demonstrates a 2.3% shift in normalized stiffness. This shift remains within the separation between materials and porosities reported in Supplementary Table 12.

porosity.

#### 2.3.2. Surrogate Material Selection

We selected surrogate materials by matching relative mechanical behavior. PLA provides a stiff, rigid response similar to PLLA [34]. TPU provides a flexible, elastomeric response similar to the superelastic behavior of Nitinol [35] (Supplementary Table 4-5).

#### 2.3.3. Compression Test Configuration

Each specimen (Figure 3d) is placed between two parallel rigid plates and compressed at a crosshead speed of 1 mm/min until full densification is reached. We test three specimens per material. Each porosity group therefore contains three overlaid curves. We record the force and crush response for each test.

### 2.4. Dimensional Analysis and Justification of Geometric Scaling

We apply the Buckingham π-theorem [36] to nine variables: seven dimensional variables and two dimensionless variables. Two fundamental dimensions yield five independent groups from the dimensional variables. We include ν and φ and obtain seven groups. We select *E* and *D* as repeating variables and obtain equation (7).

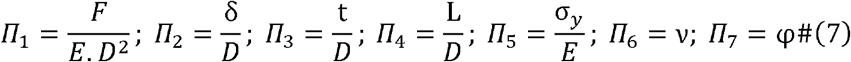

Aspect ratio similarity is exact (L/D = 1.92 in both cases). Relative strut thickness (Π3 = t/D) shows an approximately 11% deviation between the UTM specimens (0.040) and the geometrically similar target (0.045) required for exact similitude, a consequence of the minimum feature size achievable with the 0.4 mm FDM nozzle used (Supplementary Section 5).

### 2.5. Coupling Flow Rate to Radial Compliance Across the Three Stent Geometries

Because the CFD simulations assumed rigid walls, a first-order elastic-tube analysis was performed to estimate whether device radial compliance could materially alter the predicted flow. Across all nine material–porosity combinations, predicted device radial distension remained negligible relative to native vessel-wall compliance, supporting the rigid-wall assumption for the deployed-device configurations.

Using representative literature moduli, the untreated vessel wall shows a radius change of roughly 2-9% and a corresponding flow rate modulation on the order of 7-42%. These values are two to five orders of magnitude larger than those of any device configuration. The pulse wave speed relation gives a useful independent check on this. With the native wall parameters, the predicted wave speed (*c_0_* ≈ 6–14 m/s) sits within the range of *in vivo* pulse wave velocity measurements reported for human arteries [37]. Every device configuration returns *c_0_* in the hundreds to low thousands of m/s, a value with no direct physiological analogue. This result reflects how completely the device dominates the local mechanical response once deployed.

Taken together, these results demonstrate that the rigid wall simplification has negligible impact on the stented configuration. The simplification will be more consequential for the untreated, coil, and WEB models in Section 3. None of these models introduce a stiff scaffold that engages the native wall (Supplementary Section 6).

## 3. Results and discussion

### 3.1. Evaluation of Patient-Specific Model

Untreated aneurysm has a higher chance of rupture and untreated hemodynamic models of our study deliver rupture prone results. The model returns a TAWSS of 6.68 Pa, an RRT of 2.63 Pa□¹, and an OSI of 0.180, values that point to disturbed intra-aneurysmal flow presented in Figure 5.

**Figure 5.**
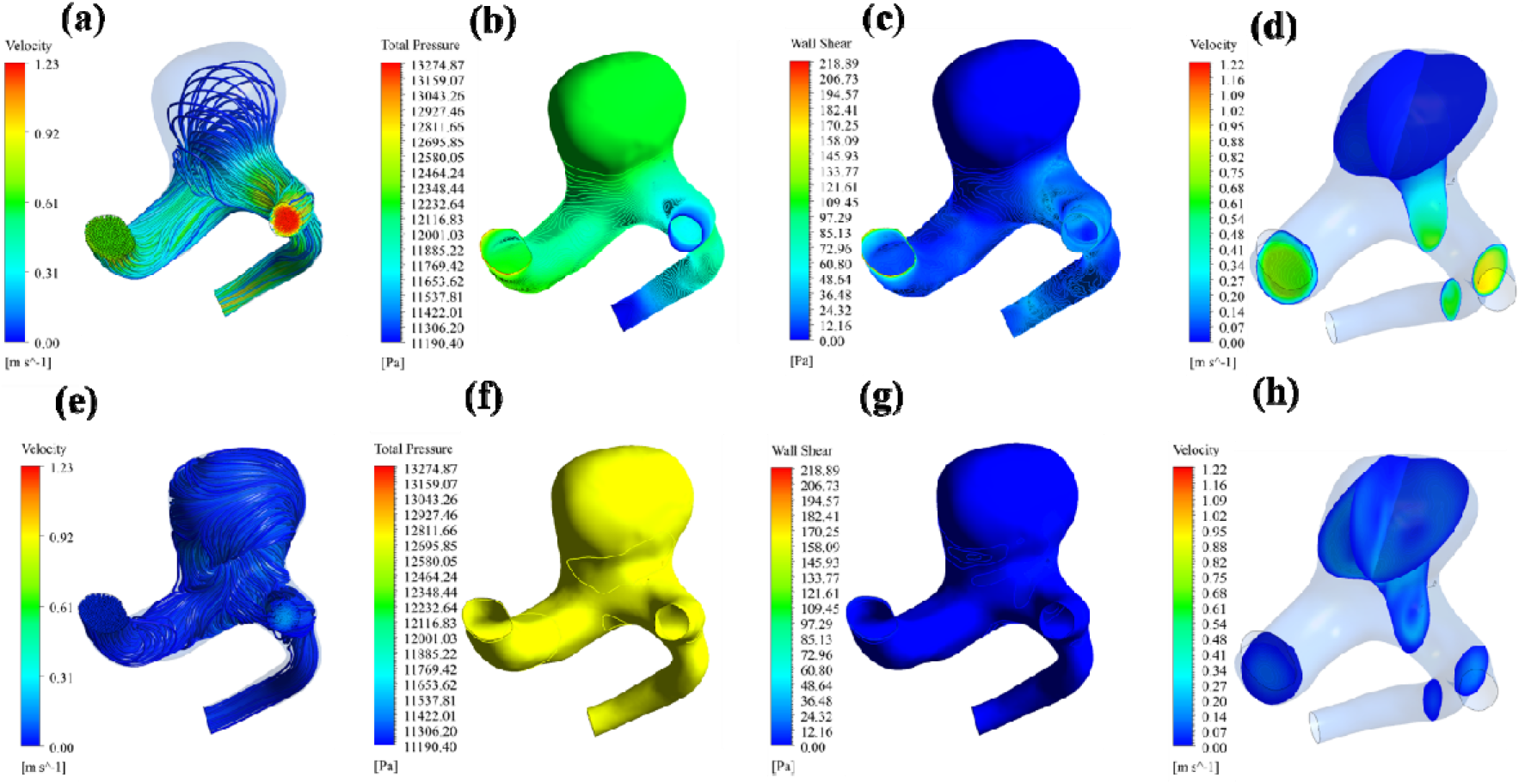
Hemodynamic characteristics of the untreated patient-specific cerebral aneurysm: (a) velocity streamlines at PS, (b) total pressure contour at PS, (c) wall shear stress (WSS) contour at PS, (d) vorticity contour at PS, (e) velocity streamlines at ED, (f) total pressure contour at ED, (g) wall shear stress (WSS) contour at ED, and (h) vorticity contour at ED.

At both peak systole (PS) and end diastole (ED), total pressure stays essentially flat across the cardiac cycle, sitting at 12061.9 Pa, which confirms stable pressure loading under the physiological conditions we prescribed. Wall shear stress (WSS) falls sharply from 29.30 Pa at PS to 2.61 Pa at ED, tracking the deceleration of blood flow during diastole. So, the aneurysm wall sees two very different shear environments depending on the position in the cycle.

Vorticity drops from 5898.44 s□¹ at PS to 153.49 s□¹ at ED. The high systolic value points to strong rotational structures forming inside the sac, and the much lower diastolic value suggests those structures dissipate as inflow weakens. The moderate OSI reflects how much the wall shear direction shifts over the cycle [38], and the elevated RRT confirms blood is sitting in the sac longer than it should [38]. Together, oscillatory shear and prolonged residence time are the hallmarks of disturbed hemodynamics [39]. These baseline values are what we compare the proposed flow diversion strategies against in the analyses that follow.

### 3.2. Hemodynamic Analysis of Flow Diversion Strategies

We report results at two extremal instants of the imposed cardiac cycle: peak systole (PS, *t* = 0.07 s) and end diastole (ED, *t* = 0.78 s). We evaluate four geometries: (a) untreated, (b) stent, (c) coil, and (d) WEB. Additionally, we calculate cycle-averaged, area-weighted wall parameters: time-averaged wall shear stress (TAWSS), relative residence time (RRT), and oscillatory shear index (OSI). Throughout, percentage changes are computed relative to the untreated baseline.

#### 3.2.1. Intra-Aneurysmal Flow Structure

Peak-systolic streamlines in the untreated model (Figure 6a) contain one coherent recirculating structure in the sac. An oblique jet impinges on the distal sac wall and loops toward the neck. Stent deployment (Figure 7b) preserves the recirculation, redistributing flow around the neck strut lattice and parent-vessel streamlines. In the coil model (Figure 6c), the densely packed wire mass leaves few continuous flow paths within the sac, whereas the parent-vessel streamlines beneath the coil mass exhibit nonuniform velocity reduction. The coil mass reduces the available flow cross-sectional area. Figure 6d presents the WEB model that demonstrates suppressed recirculation, while flow velocity remains intermediate between the stent and coil cases.

**Figure 6.**
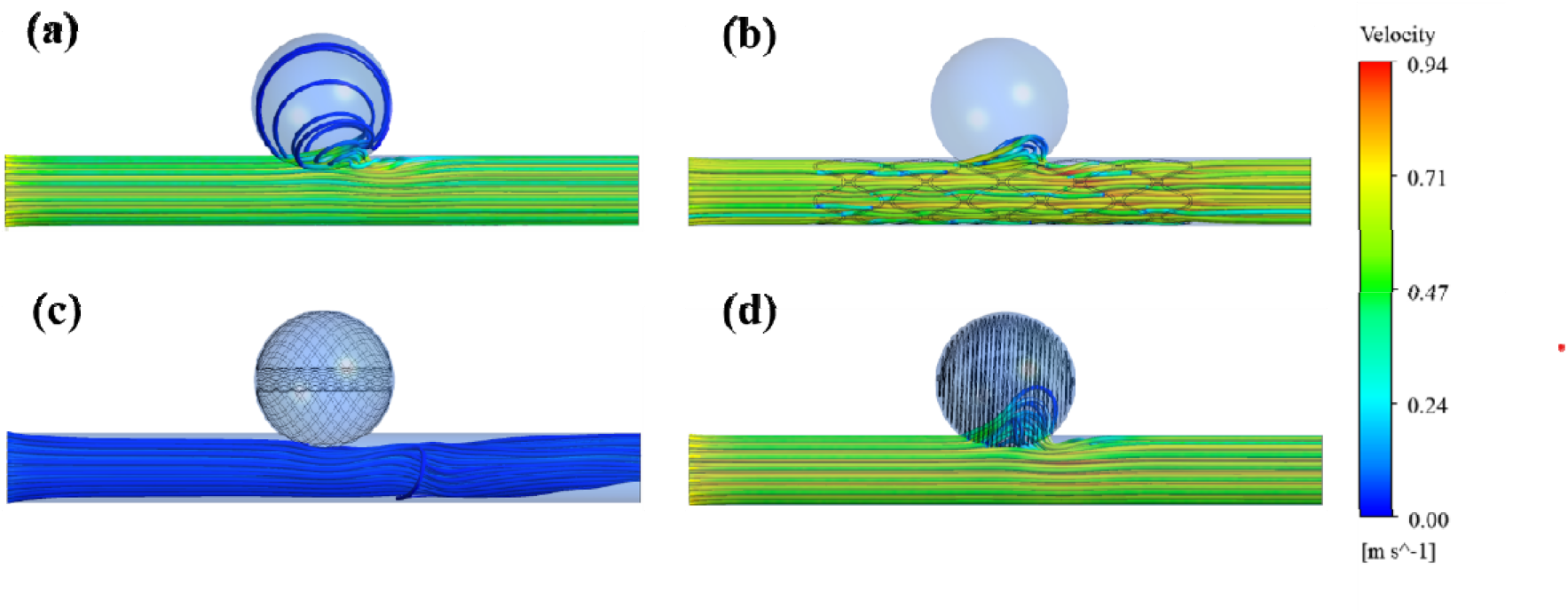
Peak-systolic (t = 0.07 s) velocity streamlines for (a) untreated, (b) stent, (c) coil, and (d) WEB models.

**Figure 7.**
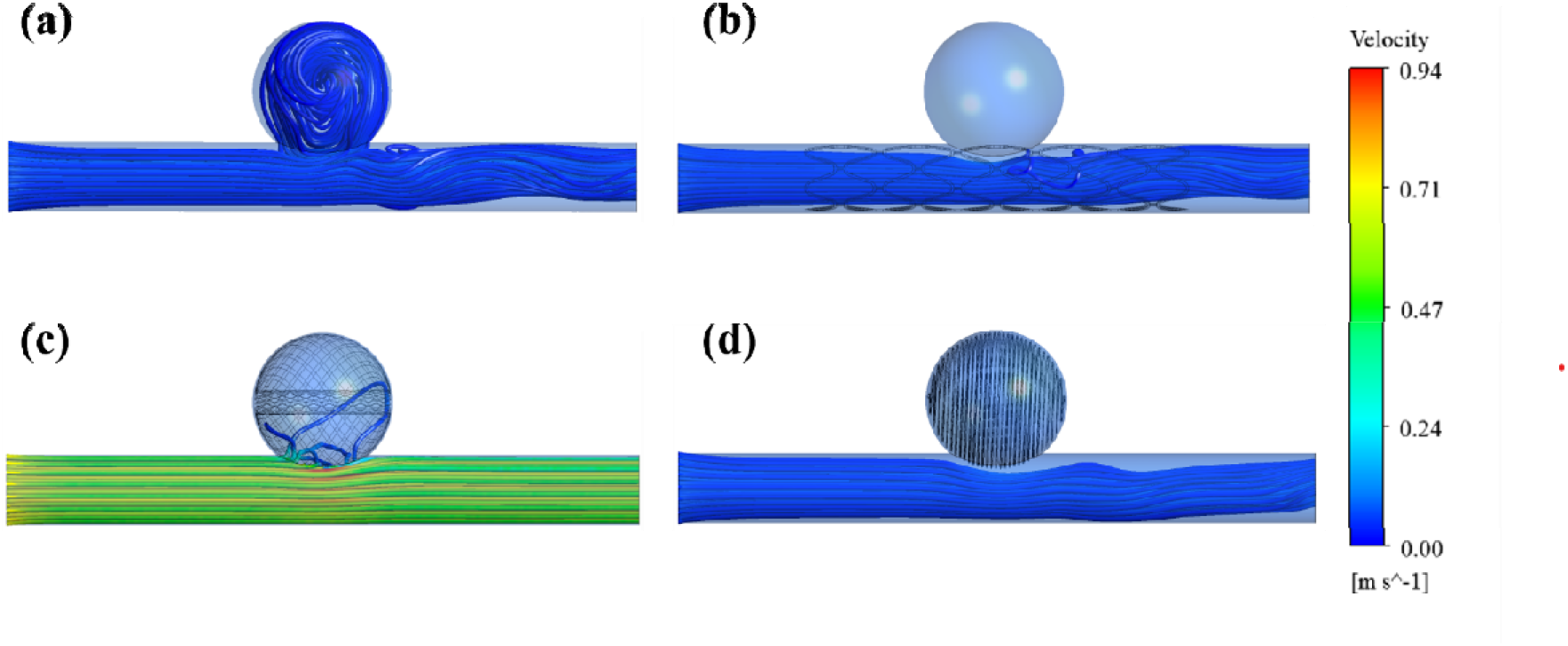
End-diastolic (t = 0.78 s) velocity streamlines for (a) untreated, (b) stent, (c) coil, and (d) WEB models.

Streamline speed drops toward the low end of the color scale across all four models, at end diastole (Figure 7), consistent with the inlet waveform at that point in the cycle. The coil model is the only one that retains noticeable velocity variation in the parent vessel, suggesting residual flow is being channeled locally around the coil mass rather than stagnating uniformly.

#### 3.2.2. Velocity Field

The peak-systolic velocity contour at the sac mid-plane (Figure 8) matches the findings got in the streamline picture. Both the untreated and stented models show large high-velocity cores spanning the sac, but the coil model flattens that out into a nearly uniform low-velocity field. The WEB model shows a smaller, spatially confined high-velocity patch.

**Figure 8.**
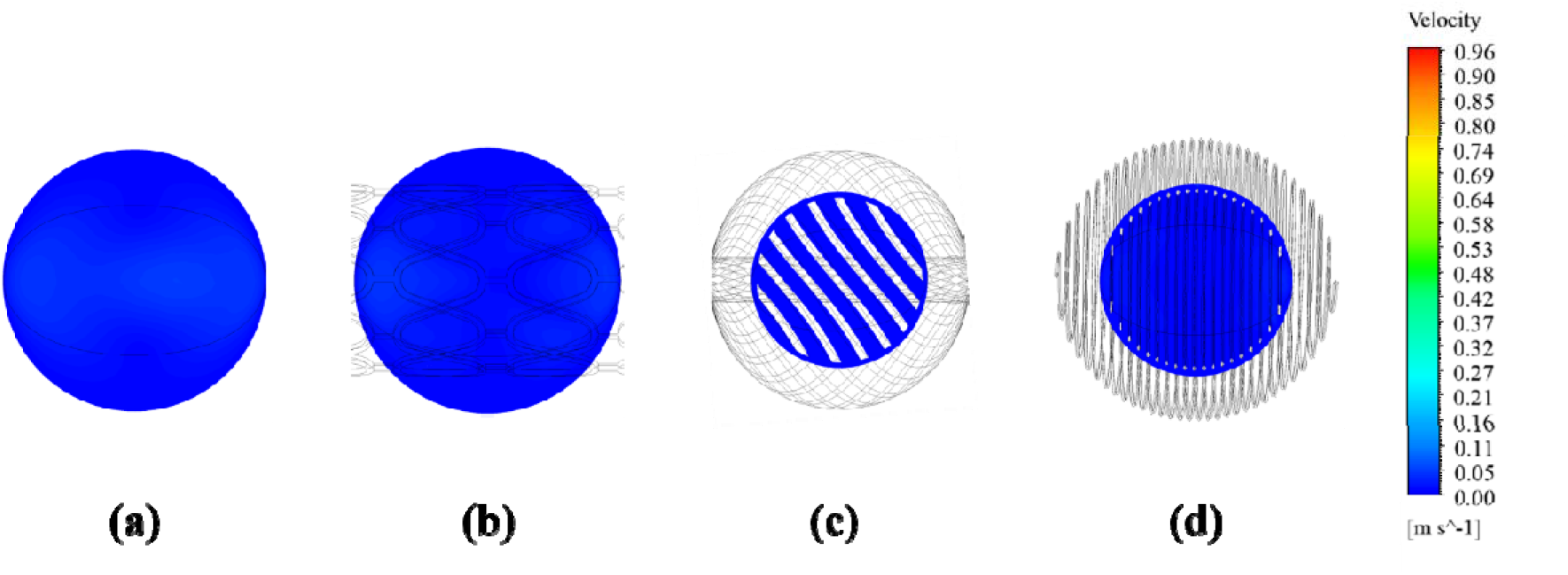
Peak-systolic velocity contour on the sac mid-plane for (a) untreated, (b) stent, (c) coil, and (d) WEB models.

The stent and WEB device cut peak systolic velocity by 16.5% and 17.3% respectively (Supplementary Table 9) relative to the untreated baseline, while the coil model actually increases it by 15.6%. That increase makes sense because the coil mass takes up a large fraction of the sac volume, so fluid squeezing through the interstitial space between loops has to locally speed up just to conserve mass flux through the narrowed cross-section.

At end diastole, all four models converge to a narrow band of 0.017 to 0.019 m/s (Supplementary Table 10), with the spread between them staying under 0.002 m/s. The stent’s 3.4% increase over baseline is real but practically meaningless at such a low-flow point in the cycle.

#### 3.2.3. Pressure Field

Static pressure was invariant across devices at peak systole and end diastole (Supplementary Table 9-10). Peak-systolic sac pressure differed from the untreated model by ≤0.6% (stent) and ≤0.3% (coil). End-diastolic pressure differed by <0.02% across all four models. The rigid-wall, incompressible-flow model enforces this invariance. The applied pulsatile pressure boundary condition directly sets the outlet static pressure.

The devices modify local velocity and shear fields more strongly than bulk pressure showing a more uniform, lower-magnitude pressure field over the sac dome in all three treated models. This pattern demonstrates inflow-jet redistribution away from the dome (Supplementary Figure 2).

#### 3.2.4. Wall Shear Stress and Vorticity

We report vorticity alongside wall shear stress (WSS). Vorticity quantifies local flow disturbance within the sac and the complex, unstable flow patterns associated with increased aneurysm rupture risk in patient-specific hemodynamic cohorts [40].

At peak systole, the stent yields the largest reductions in point WSS (-14.8%) and vorticity (-15.1%) relative to the untreated model. This result confirms the WSS reductions reported for flow-diverting stents in idealized and patient-specific computational fluid dynamics (CFD) studies [41]. The coil model increases both peak-systolic WSS and vorticity by 6.1%. The coil wires create localized high-shear, high-vorticity channels. Bulk sac flow remains disrupted. The WEB device changes peak-systolic WSS by -0.04% and vorticity by -0.08% relative to the untreated model. This dissociation between velocity and near-wall shear demonstrates that the WEB braid redirects most incoming flow away from the dome. The shear environment immediately adjacent to the device surface remains comparatively undisturbed at this instant. Figure 9 shows a low-amplitude scalloped WSS pattern in the stented parent vessel that mirrors the strut lattice. The coil and WEB sac surfaces show fine striping that follows the wire or braid orientation.

**Figure 9.**
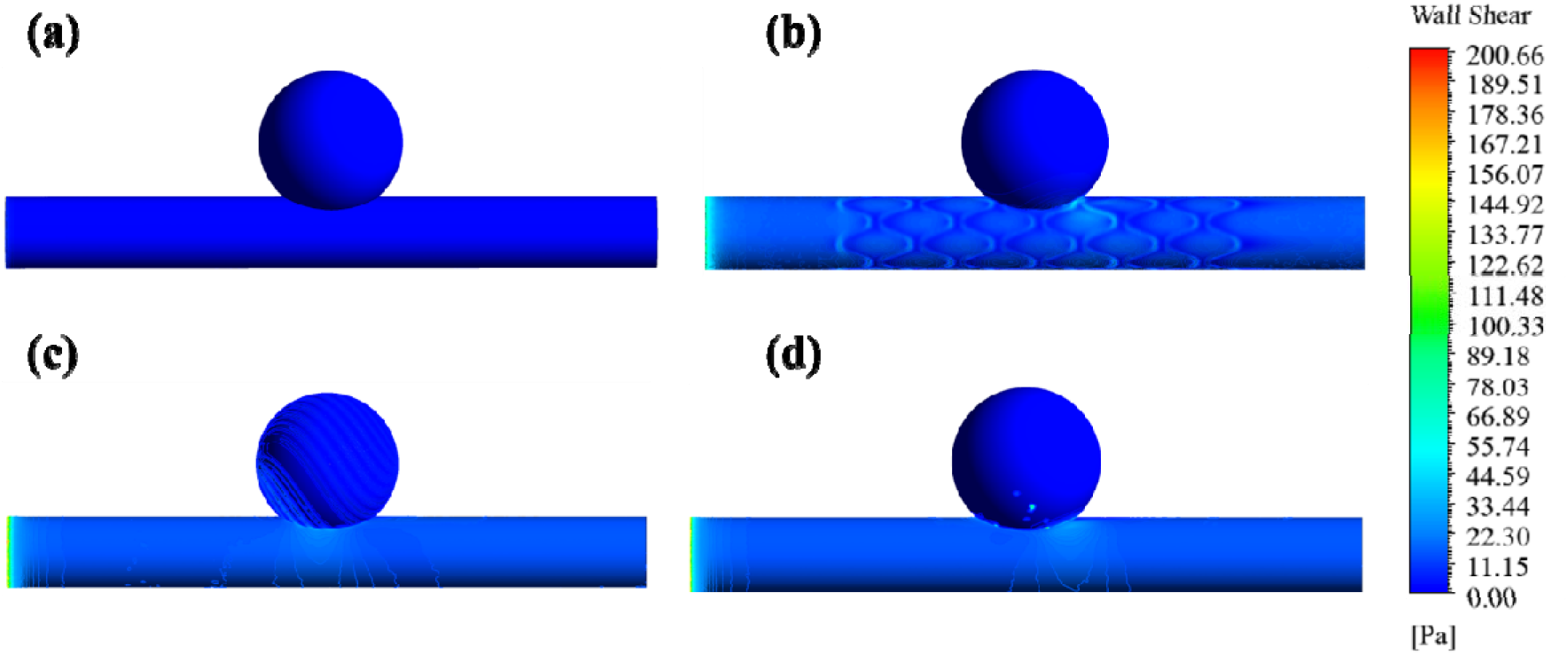
Peak-systolic wall shear stress contours for (a) untreated, (b) stent, (c) coil, and (d) WEB models.

At end diastole, the stent and coil models reduce WSS relative to the untreated model. The WEB increases WSS by 13.1%. Vorticity decreases in all treated models: coil (-18.8%), stent (-11.7%), and WEB (-2.9%). Cross-sectional vorticity fields through the parent vessel (Supplementary Figure 3) show the expected wall-adjacent shear layer in all four models with the core flow remaining close to zero vorticity in each case.

#### 3.2.5. Cycle-Averaged Wall Parameters: TAWSS, OSI and RRT

Table 2, Figures 10–11 and Supplementary Figure 4 summarize the area-weighted, cycle-averaged wall parameters. These metrics enable definitive comparison of each device’s sustained hemodynamic environment. They clearly identify the configuration most conducive to intrasaccular thrombosis [42]. Parameter change values from untreated condition is shown in Table 3.

**Figure 10.**
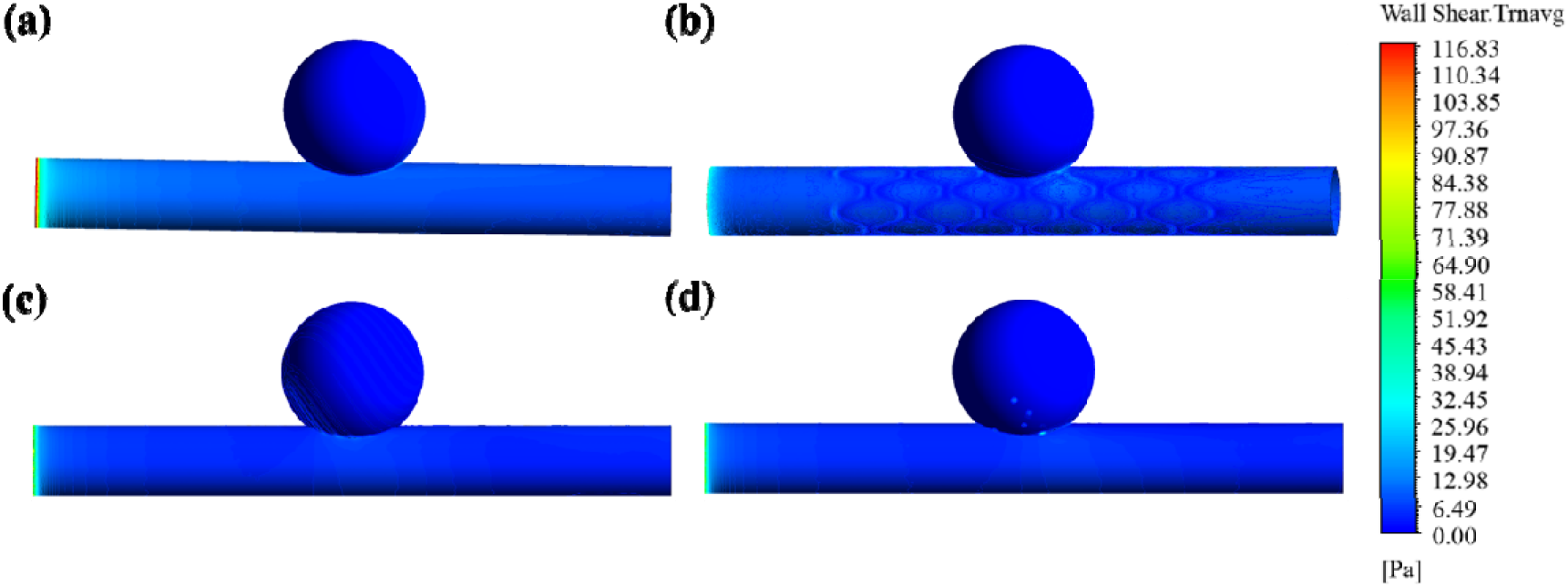
Time-averaged wall shear stress (TAWSS) distribution for (a) untreated, (b) stent, (c) coil, and (d) WEB models.

**Figure 11.**
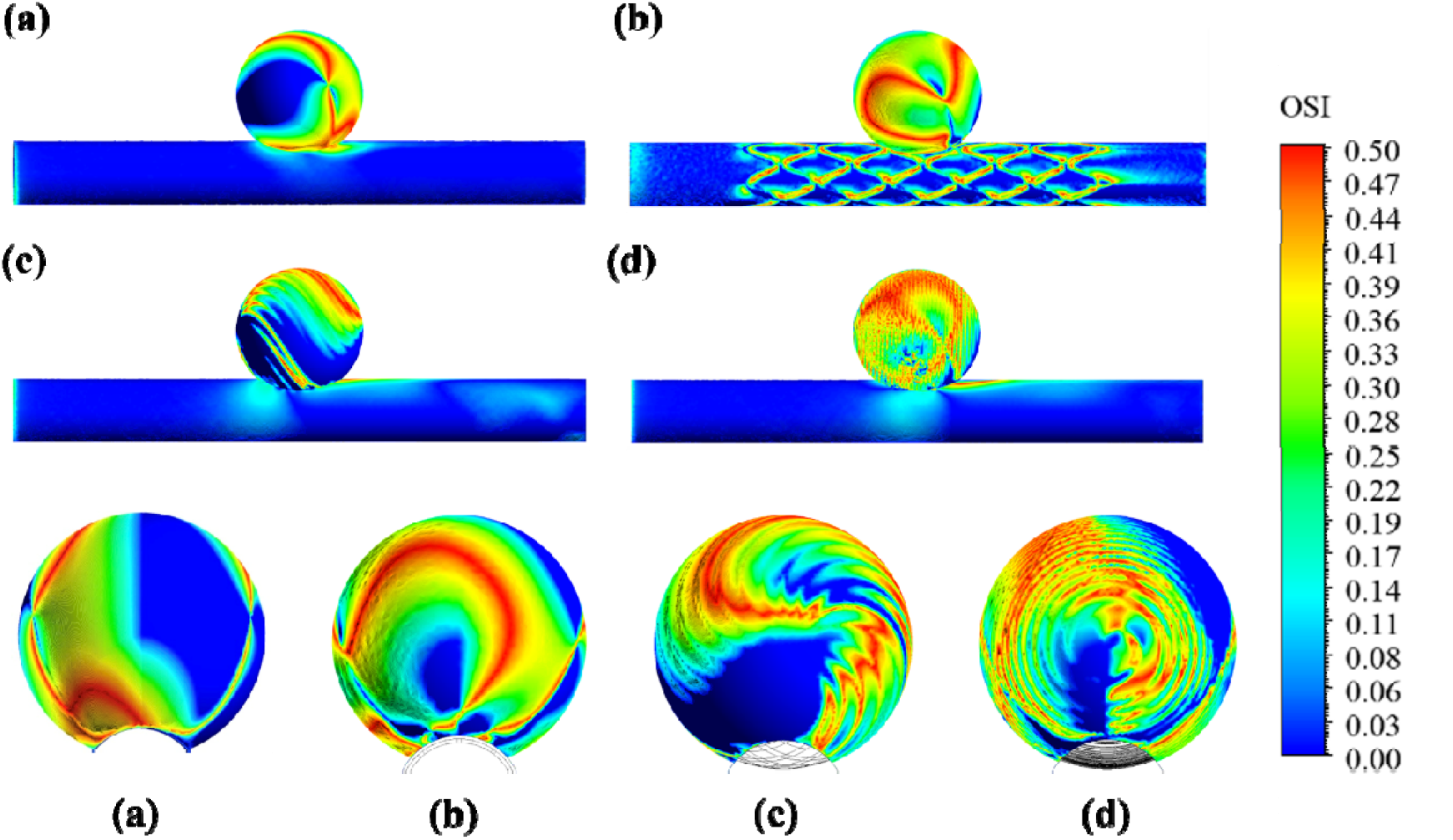
Oscillatory shear index (OSI) distribution for (a) untreated, (b) stent, (c) coil, and (d) WEB models (side and dome views).

**Table 2.** Area-weighted average hemodynamic parameters over the cardiac cycle.

| Parameter | Untreated | Stent | Coil | WEB |
| --- | --- | --- | --- | --- |
| TAWSS (Pa) | 2.856 | 2.256 | 3.133 | 2.962 |
| OSI | 0.0605 | 0.1441 | 0.0984 | 0.0832 |
| RRT (Pa<sup>-1</sup>) | 15.86 | 45.31 | 12.22 | 51.10 |

**Table 3.** Area-weighted parameters expressed as (%) change from the untreated baseline.

| Parameter | Stent ($\Delta\%$) | Coil ($\Delta\%$) | WEB ($\Delta\%$) | Direction Favoring Stasis |
| --- | --- | --- | --- | --- |
| TAWSS (Pa) | -21.0% | +9.7% | +3.7% | Decrease |
| OSI | +138.3% | +62.7% | +37.6% | Moderate increase |
| RRT (Pa<sup>-1</sup>) | +185.7% | -22.9% | +222.1% | Increase |

TAWSS decreases by 21.0% with the stent and Figure 10 presents the contour images. The stent reconstructs the parent vessel lumen and diverts flow past the neck [43]. Both coil and WEB increase area-weighted TAWSS by 9.7% and 3.7%, respectively, relative to the untreated model. The stent alone decreases this primary cycle-averaged shear metric toward the therapeutic target [44].

OSI rises in all three treated models relative to the untreated model, with an increase of 138.3% for the stent. Struts generate small, spatially periodic recirculation zones downstream. These zones locally reverse the WSS vector direction over the cycle and elevate OSI. These increases demonstrate flow diversion independent of device geometry. However, they do not offset the substantial reductions in TAWSS and peak-systolic shear achieved by the same device. OSI is not informative in isolation within a treated sac; it should be interpreted jointly with TAWSS through RRT [45]. Coil and WEB produce increases of 62.7% and 37.6%, respectively. The sac-dome OSI maps for both devices (Figure 11c–11d) show spatially periodic striping that tracks wire geometry.

RRT combines TAWSS and OSI into a single stasis index. RRT rises for the stent and the WEB (+185.7% and +222.1%, respectively) but falls for the coil (−22.9%). The coil is the only device for which area-weighted RRT (Supplementary Figure 4) drops below the untreated baseline, thereby moving this index in the therapeutically unfavorable direction. The WEB shows the numerically largest RRT increase of the three. Elevated RRT within a treated sac denotes prolonged near-wall residence time and is directly associated with the intra-aneurysmal thrombosis that underlies the therapeutic mechanism of all three devices [45].

### 3.3. Comparative Synthesis: The Stent Provides the Most Favorable Hemodynamic Profile

The stent produces the most favorable overall hemodynamic profile of the three treatment strategies modeled and Table 4 presents the overall summary with evidence, finding and interpretation. First, TAWSS is the single most widely reported cycle-averaged outcome measure in the flow-diversion CFD literature [46]. TAWSS falls only for the stent (Table 2). Both the coil and the WEB move this metric in the opposite, therapeutically undesired direction. Second, the stent produces the largest reductions in velocity, WSS, and vorticity of any device at peak systole (Supplementary Table 9). Peak systole imposes the greatest mechanical loading on the aneurysm wall. Absolute WSS and vorticity magnitudes are one to two orders of magnitude larger than those at end diastole (compare Supplementary Tables 9 and 10). Peak-systolic performance clearly favors the stent. It therefore dominates the overall hemodynamic picture, far more than the numerically small end-diastolic differences among devices.

**Table 4.**
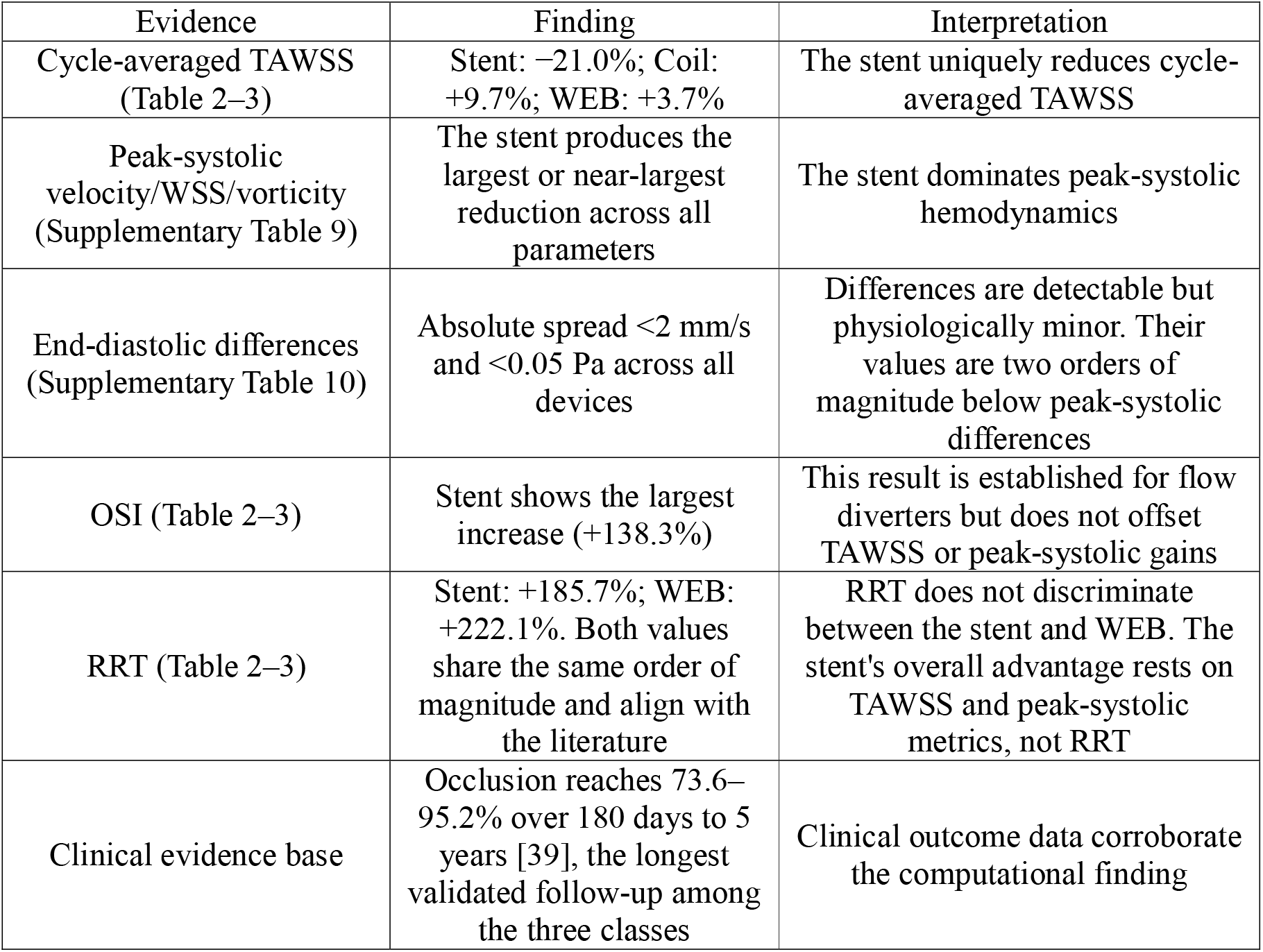
Summary of evidence supporting the stent as the preferred overall treatment strategy.

The OSI increase at the stent struts is well documented among flow-diverting stents [45,47]. It is not a deficiency specific to the modeled geometry. It accompanies the substantial TAWSS and peak-systolic shear reductions that are the stent’s principal effect. The coil is the only device that moves both TAWSS and RRT in the therapeutically unfavorable direction. Coil embolization is associated with higher clinical recanalization rates relative to flow diversion [48].

The primary cycle-averaged shear metric and instantaneous peak-systolic fields favor the stent. Weighted OSI, end-diastolic, and RRT profiles did not reduce the stent’s performance.

Clinical follow-up data corroborate this result; the stent has more extensive clinical follow-up than the WEB device and coils. The WEB device likewise establishes a stasis-favoring sac environment [49]. Unlike the stent, WEB does not reduce cycle-averaged TAWSS or dominate the peak-systolic fields. Combined computational and clinical evidence identifies the stent as the preferred flow-diversion strategy among the three modeled devices. The WEB device is a strong second-line option. The coil model has the least favorable sustained hemodynamic profile and a higher documented clinical recanalization rate [50].

### 3.4. Finite Element Analysis

Across all three materials, reaction force and radial stiffness decreased consistently with increasing porosity. For 316L SS, stiffness dropped from 45.22 N/mm at 50% porosity to 16.27 N/mm at 60%; a similar trend held for Nitinol (14.71 to 9.67 N/mm) and PLLA (0.88 to 0.58 N/mm). This confirms that reducing strut cross-sectional area lowers the structure’s resistance to radial compression regardless of material (Supplementary Table 11).

All three materials showed a dip at 55% porosity. Values rose again at 60% rather than decreasing steadily. This reflects a transition between two competing strut-level mechanisms rather than a numerical artifact. Denser strut packing reduces inter-strut spacing at crown regions at 50% porosity. This concentrates stress at geometric discontinuities. Fewer load-bearing struts force each remaining strut to carry a proportionally larger share of the applied load at 60% porosity. Section 3.9 develops this distinction further as part of the porosity-performance regime map (Figure 17b).

Material comparison demonstrated that, at every porosity level, 316L SS exhibited the highest radial stiffness. Then the other two materials, Nitinol showed considerably larger equivalent strain values. These values were an order of magnitude higher in most cases, reflecting stress-induced transformation behavior rather than simple elastic deformation. PLLA exhibited the lowest stress magnitudes and stiffness values, consistent with its lower modulus, though its strain levels remained comparable to Nitinol’s.

### 3.5. Force-Displacement Response, Radial Strength, and Stiffness

Across all nine combinations, reaction force rose nonlinearly with displacement, but the shape of that response depended heavily on material (Figure 12). 316L SS was steeply at first, then flattened into a plateau before steepening again near maximum compression, showing a pattern that points to the elastic-to-plastic transition kicking in along with strut-contact effects. Nitinol and PLLA both increased smoothly and nearly linearly across the full displacement range, and this held true at all three porosities.

**Figure 12.**
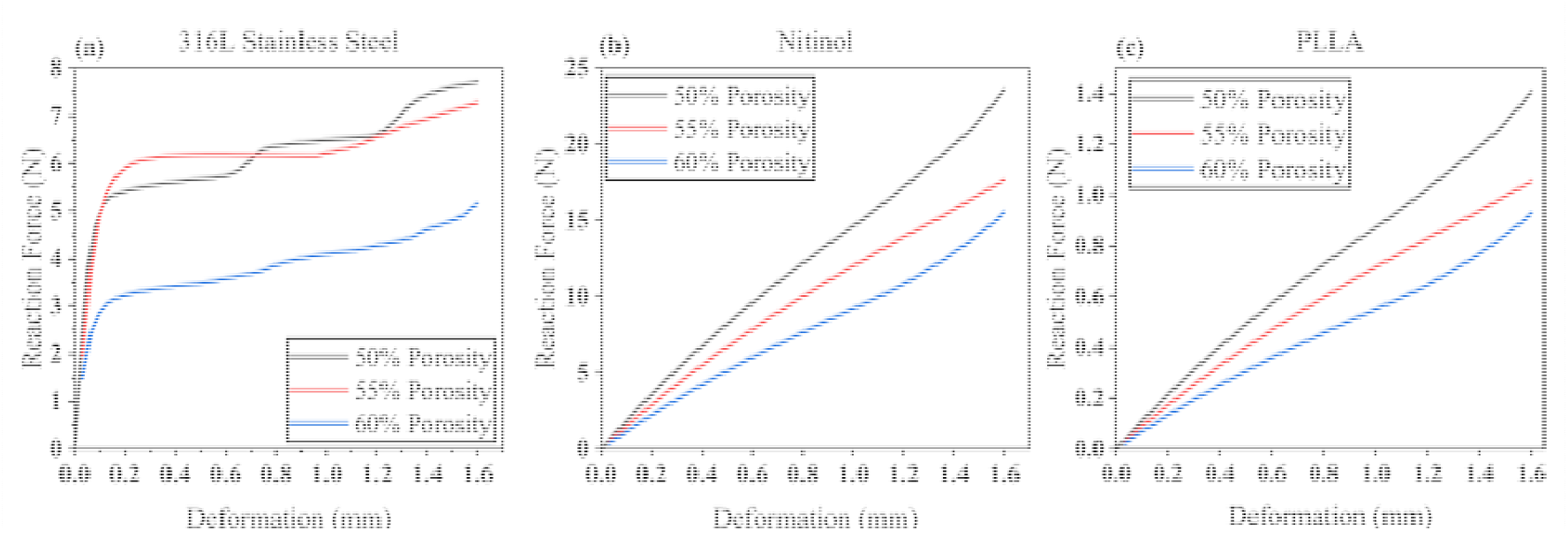
Reaction force versus displacement curves during parallel-plate compression for (a) 316L SS, (b) Nitinol, and (c) PLLA at 50%, 55%, and 60% porosity.

Because of that difference in behavior, we calculated stiffness differently for each material. For 316L SS, we used the 0 to 0.1 mm slope, since that’s the window before plastic yielding starts bending the curve. For Nitinol and PLLA, where the response stayed linear throughout, we used the full 0 to 1.6 mm slope instead.

Figure 13 summarizes radial strength and stiffness across all cases. Nitinol produced the highest radial strength at every porosity level. Its strength was approximately three times that of 316L SS and exceeded that of PLLA. 316L SS produced the highest radial stiffness, exceeding Nitinol and PLLA at 50% porosity. Both metrics decreased as porosity increased in all three materials. PLLA showed the lowest values for both strength and stiffness throughout, reflecting its low elastic modulus. The 50% porosity 316L SS design provided the greatest combined resistance to radial compression. The 50% porosity Nitinol design provided the highest load-bearing capacity with greater compliance.

**Figure 13.**
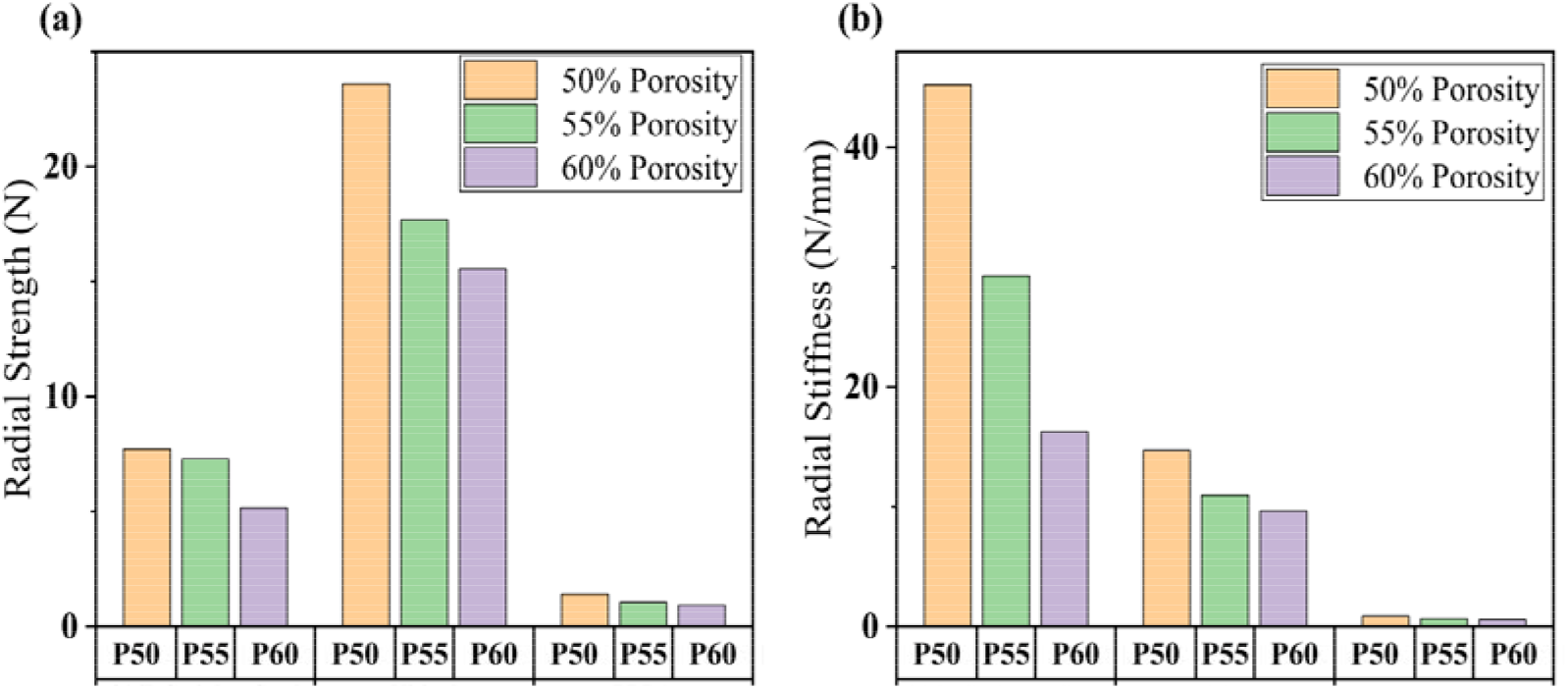
Comparison of (a) radial strength and (b) radial stiffness across all material and porosity combinations under parallel-plate compression.

### 3.6. Equivalent Stress (Von-Mises stress) and Equivalent Strain

Equivalent stress and strain distributions pinpoint where each design peaks under radial compression, and in all nine cases that turned out to be the same place: strut junctions and crown regions. Unlike the straight strut segments, those spots bend severely during compression.

Figure 14 shows the stress and strain contours for all three materials at all three porosities, with stress on the left (a-i) and strain on the right (j-r). 316L SS concentrates its highest stress into small, localized red spots at the crown tips, matching where plastic yielding is expected to start. Nitinol spreads elevated stress and strain more broadly across the strut network instead of sharply localizing it, which fits with stress-induced phase transformation. Larger regions of material transform together rather than a single point yielding first. PLLA’s stress pattern looks visually similar to 316L SS, concentrated at the crowns, but its strain values run an order of magnitude higher, which tracks with its much lower elastic modulus.

**Figure 14.**
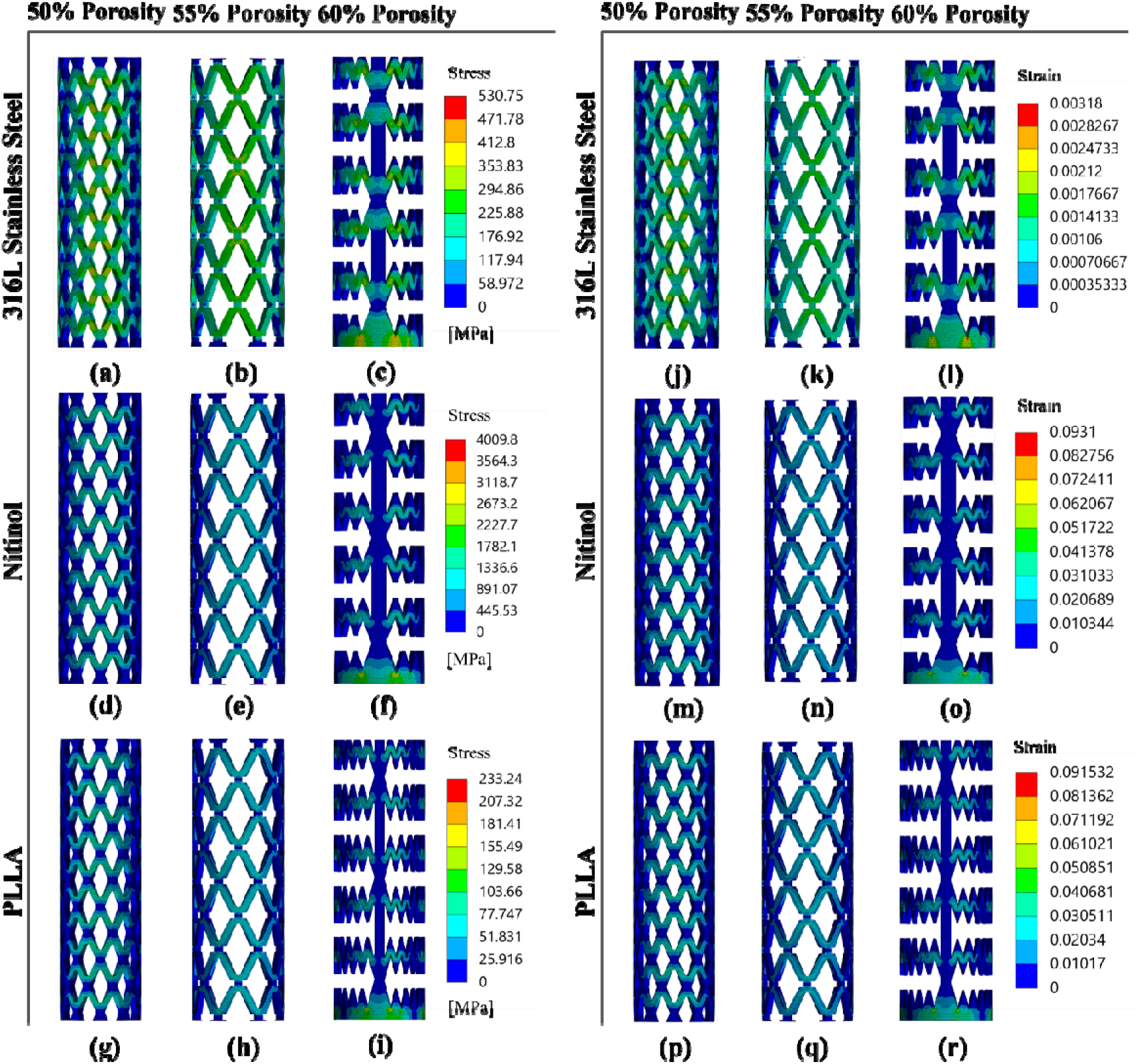
Equivalent stress (a to i) and equivalent strain (j to r) contours at maximum compression for 316L SS, Nitinol and PLLA at 50%, 55%, and 60% porosity.

Peak stress and strain didn’t scale predictably with porosity. We didn’t see a monotonic increase in either metric (Supplementary Table 11), going from 50% to 60% porosity across all three materials. Instead, the 55% configuration consistently came out lowest on both counts, which suggests strut geometry and junction spacing at that porosity happen to distribute load more evenly than at 50% or 60%.

### 3.7. Experimental Compression Behavior

Figure 15 shows compression responses for polylactic acid (PLA) and thermoplastic polyurethane (TPU), with three overlaid curves per material at each porosity group. Both materials traced the same three-phase response typical of porous cellular structures [51]: an initial linear region where the struts deform elastically, a plateau where the strut network progressively collapses at roughly constant force, and a densification region where force spikes once the collapsed struts start contacting each other.

**Figure 15.**
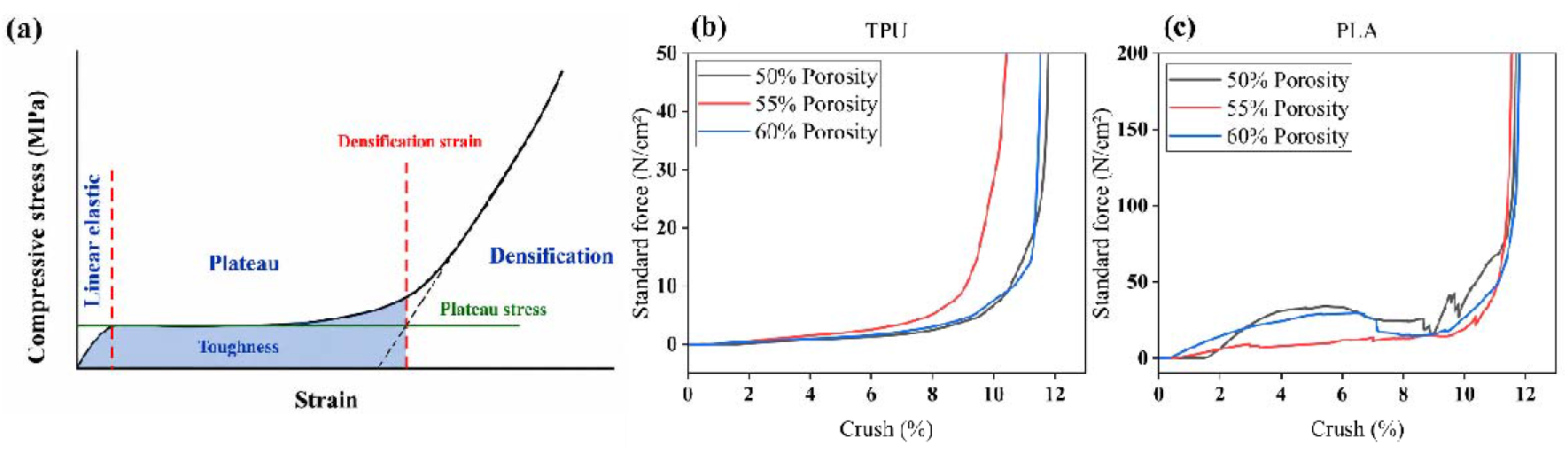
(a) Generalized Curve of Compression Test in Porous or Cellular Structure [52] and Compression Test Curve of (b) TPU, and (c) PLA.

The TPU specimens, representing nitinol, showed a longer, more gradual plateau before densification. This response demonstrates a more flexible, elastically recoverable structure. The PLA specimens, representing PLLA, showed a shorter plateau and an earlier onset of densification, indicating a stiffer, more brittle response. At 50% porosity, specimens entered densification at a higher force and later in the crush range than at 55% or 60% porosity. Lower-porosity designs therefore provide greater resistance to compression before structural collapse.

The 50% porosity design demonstrated the highest this property for both materials. It maximized force retention through the plateau region and delayed densification. TPU (representing Nitinol) exhibited gradual, controlled collapse. PLA (representing PLLA) showed a stiffer initial response and collapsed more abruptly after the plateau ended. This ranking confirms the porosity trend observed in the simulated radial strength and stiffness results.

### 3.8. Dimensional Scaling Verification

We tested geometric scaling between FEA models (4 mm diameter) and UTM specimens (25 mm diameter) for mechanical equivalence. We computed normalized stiffness Π_1_ = k/(E·D) (Section 2.4) for 9 FEA combinations and 6 UTM combinations. We used the corresponding measured or literature elastic moduli. Supplementary Table 12 reports Π_1_ for the FEA dataset. It also reports analogous UTM values computed from measured PLA and TPU force–displacement slopes and their respective moduli. We extracted elastic modulus (E) values from Section 2.2.2. We calculated the *k* values for PLA and TPU from their slopes. We converted crush percentage to displacement (δ = crush% × 48 mm). We used the initial linear region of each curve.

Figure 16 shows that, FEA and UTM datasets exhibit the same porosity-dependent trend, consistent with geometric similarity (Π_3_, Π_4_ matched). FEA and UTM use different material parameters (ν, σ_y_/E). UTM uses polymer surrogates rather than the true stent materials, producing the residual absolute Π_1_offset.

**Figure 16.**
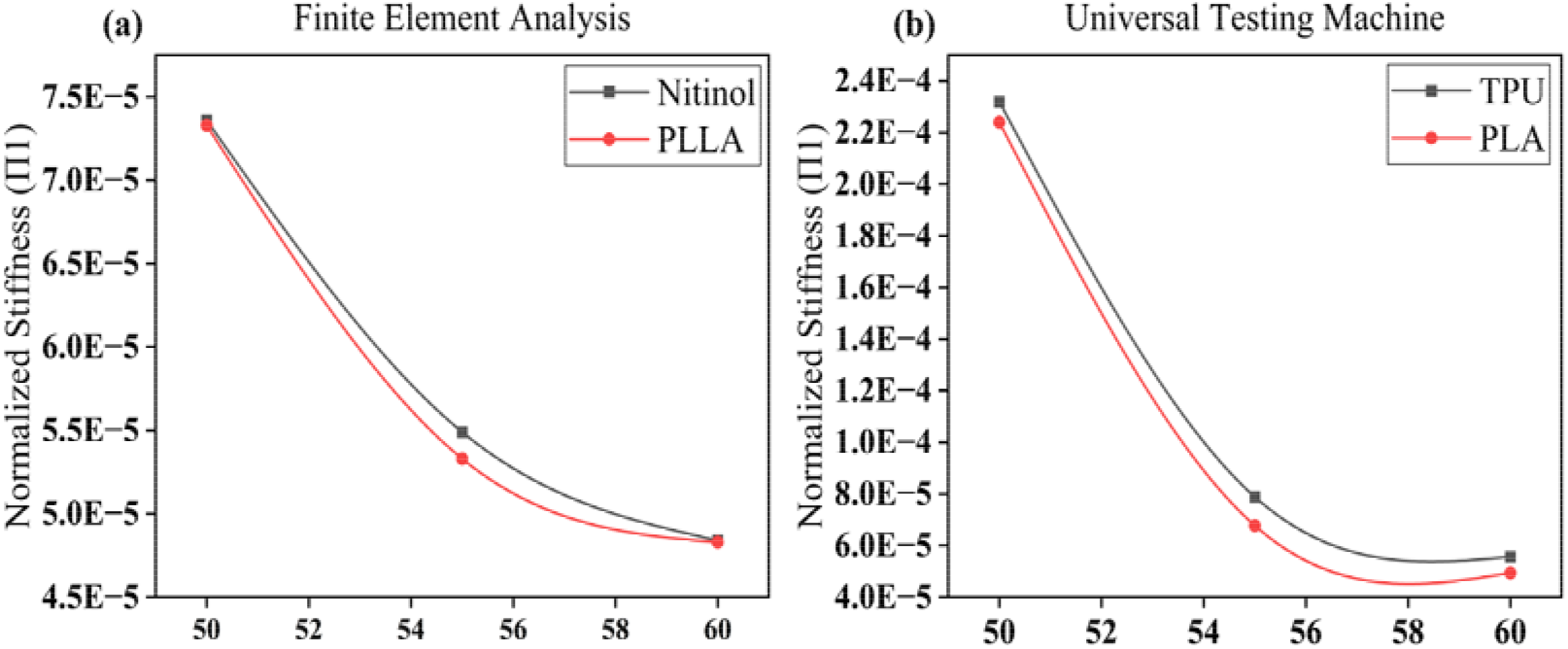
Π_1_ against porosity for the (a) FEA overlaying Nitinol, and PLLA (b) UTM overlaying TPU and PLA.

Two supplementary FEA cases used t/D = 0.040. This value matches the strut-thickness ratio realized in the printed UTM specimens, not the design value of t/D = 0.045. At fixed L/D and porosity, these cases produced a 2.3% Π_1_ shift relative to the design-geometry FEA cases. The 2.3% value is based on the 60% porosity PLLA case. This minimizes the material-surrogate effect and isolates the geometric scaling deviation. This quantifies the scaling deviation introduced by the 0.4 mm FDM nozzle resolution limit reported in Section 3.10. The UTM-based stiffness rankings demonstrate accuracy within 2.3% of the exact-geometric-similarity value. This deviation remains below the separation observed between materials and porosities in Supplementary Table 12.

### 3.9. Development of a Porosity-Performance Design Regime Map

This study evaluated three discrete porosity levels (50%, 55%, and 60%) using finite element analysis. These results establish a preliminary porosity-performance design framework for flow-diverter stents. In practical stent development, porosity is a fundamental design variable [53]. We therefore estimate structural response at intermediate porosities to reduce computational iterations during preliminary design optimization.

We propose a porosity-performance design regime map based on the mechanical behavior we observed across the finite element simulations. The idea is to treat porosity as the main variable governing shifts in structural performance. This map is meant as a starting point, not a replacement for finite element analysis; engineers can use it to narrow down candidate porosities before running the detailed numerical verification themselves.

As porosity increases, radial stiffness and reaction force both decrease consistently. This held across every material we tested. Lower porosity means a larger load-bearing structural network stays intact, so radial-compression resistance goes up, while higher porosity leaves less structural material to resist deformation, which pulls radial support back down. So, the relationship with porosity is clean and monotonic on the global stiffness front.

Across all three materials, both maximum equivalent stress and equivalent strain bottom out at 55% porosity rather than trending in one direction. This comes down to how porosity fraction interacts with strut arrangement and junction geometry to concentrate stress within the stent [54]. The lattice is denser and overall stiffness goes up at 50% porosity, but that density also produces sharper local stress concentrations. At 60%, there aren’t as many struts left to carry the load. So, the stresses that do occur get pushed through fewer paths and end up larger. Load redistribution works in 55% configuration and peak structural demand drops.

Based on this, we sorted the stent designs into three design regimes, that is shown in Figure 17. We normalized each material’s data to its own 50% porosity value, the bold curve with the shaded band shows the mean trend and range across 316L stainless steel, Nitinol, and PLLA. The background shading marks the three regimes we’re proposing. Region 1 is for high radial support, Region 2 is for balanced performance, and Region 3 is for high flexibility.

**Figure 17.**
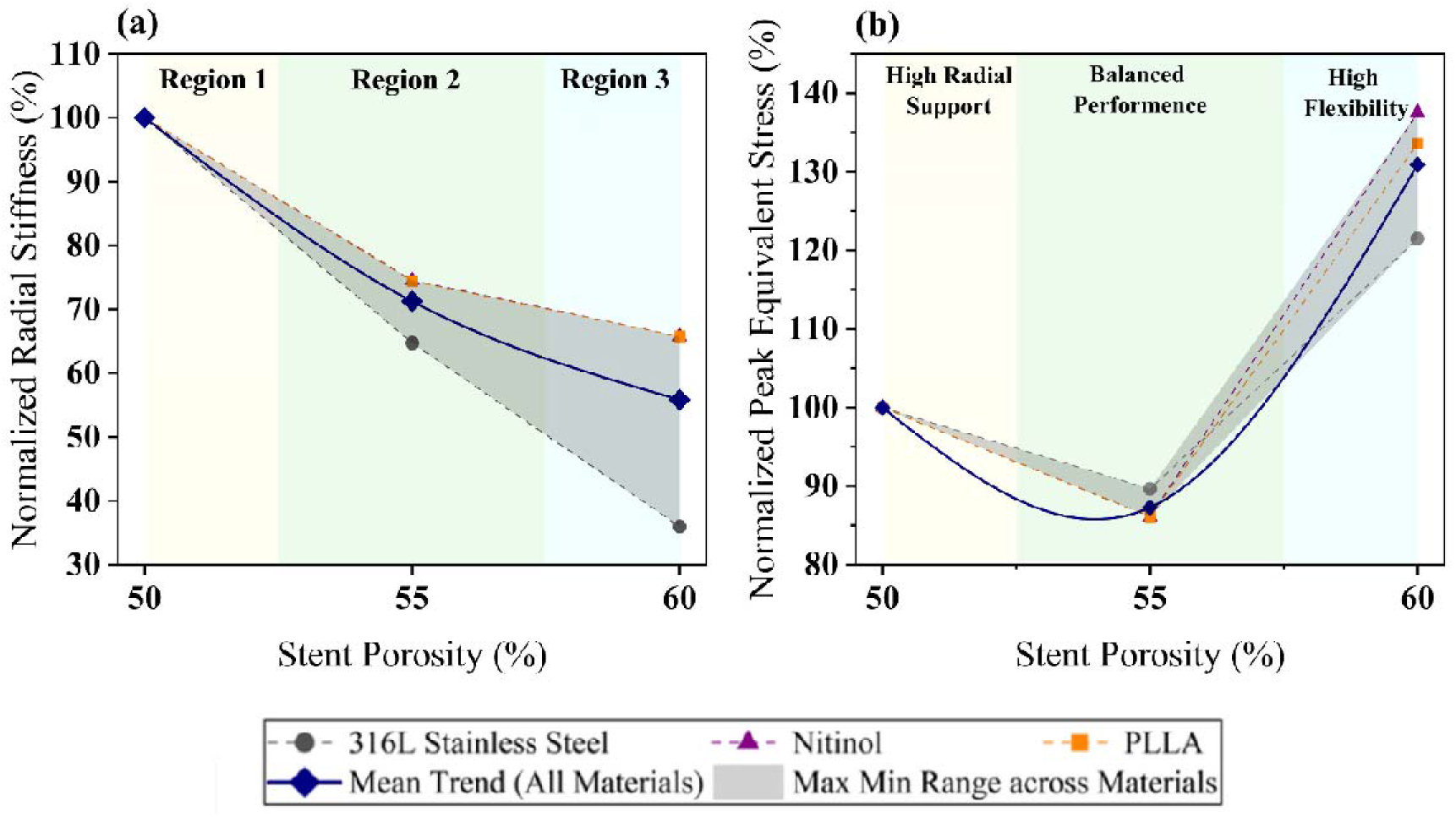
Porosity-performance design regime map for (a) normalized radial stiffness and (b) normalized maximum equivalent stress versus stent porosity.

Region 1 covers the lower-porosity designs, including the 50% configuration we tested here. These stents deliver the highest radial stiffness and reaction force, making them best suited for resisting radial compression and scaffolding the vessel wall, though the denser strut network also means higher local stress concentrations.

In the 55% configuration, the radial stiffness drops compared to the 50% design, but equivalent stress and strain hit their minimum values across all three materials. That combination points to better load redistribution and a more uniform stress field overall, which is why we consider 55% the most mechanically balanced option of the ones we tested.

At the high end, 60% porosity designs show the lowest radial stiffness and reaction force. That trade-off buys more structural flexibility and conformability, but it comes at a cost. Fewer load-bearing members means local stress concentrations go up while overall resistance to radial deformation goes down.

We see this regime map as a preliminary design tool rather than a finished one. In practice, engineers would start by deciding what mechanical response they actually want, whether that’s stiffness or flexibility, and use the map to identify a rough porosity target before moving into detailed finite element analysis. This map only covers the porosity range we actually tested, 50% to 60%, so it shouldn’t be stretched beyond those bounds. Filling in more finite element simulations at intermediate porosities would let us build out continuous empirical relationships and pin down the regime boundaries more precisely, turning this from a qualitative sketch into something closer to a predictive tool for optimization work.

The proposed porosity-performance regime map generalizes the structural findings beyond three discrete geometries. It provides an engineering framework for stent optimization. The framework does not identify a single optimum porosity. It links porosity selection to the required balance among radial support, structural integrity, and flexibility for each clinical application.

### 3.10. Comparison Among Methods to Study Aneurysm Hemodynamics and Device Structural Behavior

Most prior work isolates a single device class, a single evaluation modality, or a single data source. CFD studies focusing stent studies quantify oversizing and porosity effects but do not report any structural response [28,41]. Coil studies quantify packing density and hemodynamics but do not extend to stents or WEB devices [15,19]. The WEB literature remains hemodynamic and single-device [49]. Only structural work characterizes stent mechanics limited to a single material under compression, with experimental validation [54]. Fluid Structure Interaction (FSI) studies establish rigid-wall assumptions overestimate wall shear stress but do not compare device classes [12]. In vitro comparisons validate flow-diverter geometry experimentally but without patient-specific anatomy, finite element characterization [24]. Reproducibility studies confirm that CFD outputs are sensitive to modeling choices across repeated analysis of the same aneurysm, reinforcing the need for methodological transparency [46]. Table 5 shows the present framework against representative computational and experimental studies of aneurysm treatment published within the past decade.

**Table 5.**
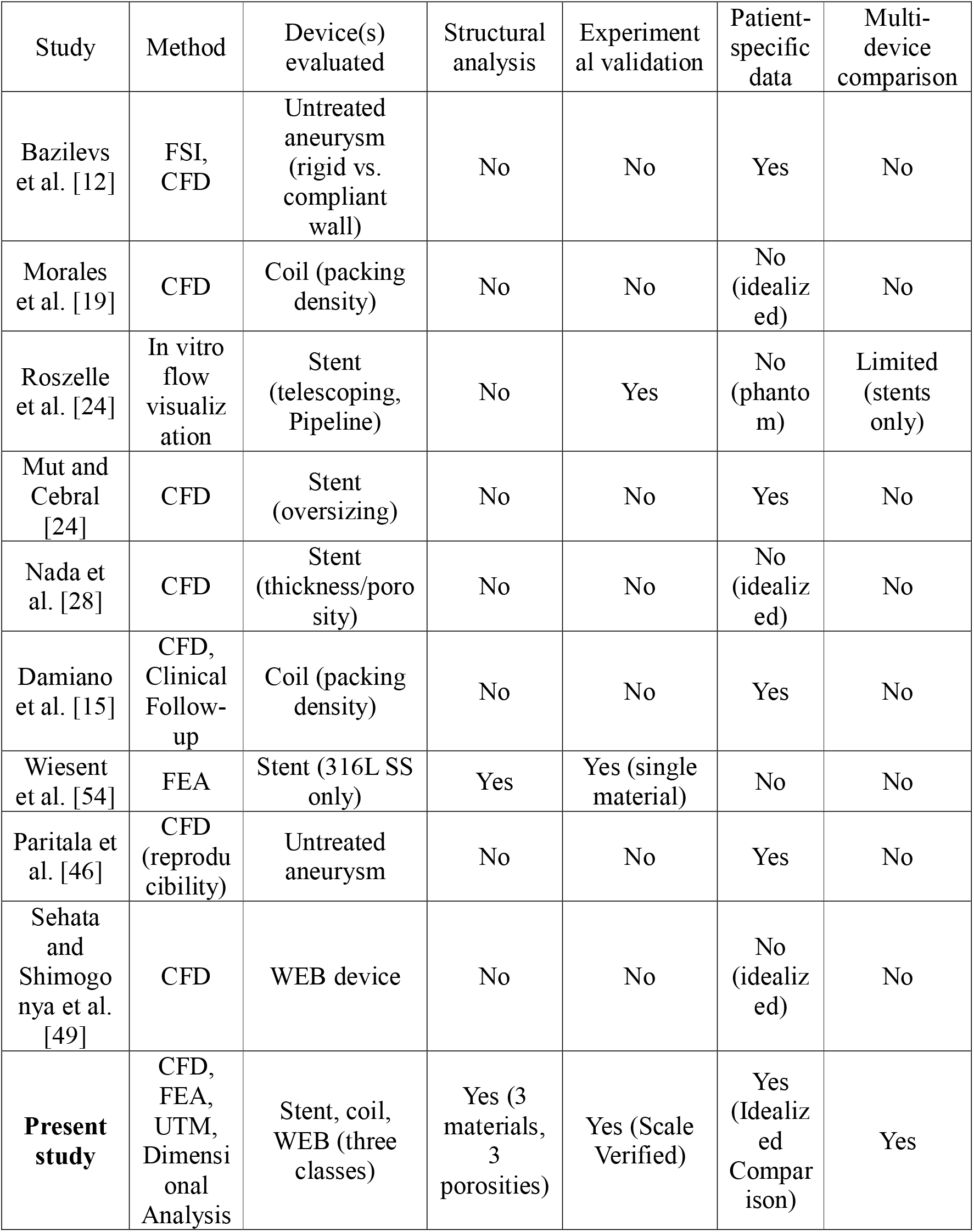
Comparison of recent computational and experimental studies of cerebral aneurysm hemodynamics and endovascular device structural behavior, contrasted with the integrated hemodynamic-structural-experimental framework applied in the present study.

The present study differs from each of these by combining four elements within one framework. Hemodynamic comparison across all three principal device classes under matched boundary conditions, structural finite element analysis across three materials, three porosities, scaled experimental validation with dimensional-analysis justification, and a closed-form check of the rigid-wall assumption. No single study in Table 5 reports more than two of these four elements together. The comparison also reflects a South Asian patient-specific data, a population underrepresented in the reviewed cohorts.

### 3.11. Limitations, Scope and Future Directions

The modeling framework limits generalization across four domains: hemodynamics, structure, fabrication, and translation.

First, all four device-comparison models treated the parent vessel and implanted devices as rigid, no-slip boundaries. We therefore interpret the reported flow, pressure, and shear magnitudes as device rankings under a matched rigid-wall assumption, not as absolute values. Fluid-structure interaction (FSI) studies of intracranial aneurysms consistently demonstrate that rigid-wall models overestimate wall shear stress and systolic velocity relative to compliant-wall simulations [12]. Relative residence time (RRT) and oscillatory shear index (OSI) are the most sensitive parameters evaluated here. Their magnitudes and rank order among the stent, coil, and WEB could shift under compliant-wall modeling. The TAWSS-based conclusion favoring the stent could persist. The elastic-tube analysis in Section 2.5 quantifies this concern for the deployed devices. Predicted radial distension across all nine finite-element analysis (FEA) configurations is several orders of magnitude smaller than that of the native vessel wall. This result supports the rigid-wall assumption for the stented segment. The same cannot be assumed for the untreated, coil, and WEB models, none of which introduce a stiff circumferential scaffold, and compliant-wall FSI remains the more consequential open question for those three configurations. Second, the four-way device comparison used a single idealized sidewall geometry parameterized from literature-typical dimensions, while we used the patient-specific reconstruction only to confirm baseline rupture-risk status, not for device comparison; aneurysm morphology is itself a recognized determinant of the hemodynamic environment [22]. Third, each device was represented by one configuration: one coil packing arrangement, one WEB size, and one stent design. Each configuration was evaluated over a single cardiac cycle at a fixed heart rate. Clinical coiling outcomes are known to be packing-density-dependent [55] and are achieved through staged framing, filling, and finishing coils rather than the single helical coil modeled here [56]; therefore, the comparatively unfavorable hemodynamic signature attributed to the coil does not necessarily extend across the full range of clinically achievable packing densities.

The elastic-tube coupling introduced in Section 2.5 is a first-order, quasi-steady approximation. The Womersley number for this cardiac cycle (α = 0.8–3.1) falls between the quasi-steady and inertia-dominated limits. The resulting flow-rate sensitivity therefore provides a comparative, order-of-magnitude estimate, not a phase-resolved prediction. The native-wall reference values were obtained from the literature. We homogenized the porous-stent lattice modulus using the as-designed strut thickness, not a patient-specific or manufacturer-specified shell equivalent. A full fluid-structure interaction simulation is required to resolve these effects quantitatively.

The finite element analysis (FEA) component was limited to a single, monotonic quasi-static compression to 1.6 mm. We did not simulate cyclic or fatigue loading. Repeated *in vivo* pulsatile loading motivated this study. Additively manufactured polymers also exhibit fatigue lives that differ substantially from static-strength predictions [56].

The 316L stainless steel results require a specific caveat. The alloy was modeled as linear elastic, yet the model computed peak equivalent stresses at 50%, 55%, and 60% porosity. A real strut would yield and redistribute load before reaching these stress levels. Its force-displacement plateau instead reflects large-deflection geometric softening of the strut lattice, not genuine material plasticity. The absolute steel stress values therefore quantify relative porosity effects, not literal stress predictions. Nitinol and PLLA were modeled as linear elastic and therefore did not capture their true nonlinear behavior. We simplified Nitinol’s superelastic phase transformation and PLLA’s viscoelastic response for comparison. This simplification may reduce prediction accuracy. Future work should implement their constitutive models. The three porosity geometries isolated porosity and material effects. They did not replicate the braid angle or cell geometry of a named commercial device. Parallel-plate compression approximates, but does not reproduce, the distributed circumferential loading that a stent experiences against the vessel wall *in vivo* [57].

Fabrication constraints prevented manufacturing 316L SS stents matching the simulated porosities for experimental testing. We therefore performed compression testing on Nitinol and PLLA surrogate specimens at all three porosities (n = 6). We could not print the FEA stent geometry at that scale, so we enlarged it to 48 mm length, 25 mm diameter, and 1 mm struts for UTM testing. This preserved the length-to-diameter ratio exactly but did not preserve the strut-thickness-to-diameter ratio. Geometric similarity required a 1.125 mm strut at this scale. The printed 1 mm strut caused an approximately 11% relative deviation because of printer-resolution limits. Section 3.8 quantifies the mechanical consequence of this departure directly. Supplementary FEA cases run at the as-printed strut ratio show a normalized stiffness shift of approximately 2.3%, well within the separation observed between materials and porosities (Supplementary Table 12). The UTM data therefore provide scale-verified validation of the porosity ranking and material contrast established by FEA. The surrogate materials compound this gap. TPU’s elastic modulus is roughly three orders of magnitude below Nitinol’s [58]. TPU reproduces Nitinol’s qualitative gradual recovery character but cannot reproduce the stress-induced martensitic transformation that gives clinical Nitinol its particular combination of strength and superelastic recovery [59]. A related limitation is that the FEA simulations were compressed only to 40% (1.6 mm), while the UTM specimens were compressed fully into densification, limiting direct comparison to the range covered by the simulations.

WSS, TAWSS, OSI, and RRT served as established hemodynamic surrogates for thrombogenic potential [11]. Simulations were run for slightly more than one cardiac cycle from an impulsive start rather than multiple cycles to a verified periodic state, so a residual start-up transient cannot be ruled out in the reported instantaneous and cycle-averaged values. Thrombus formation, endothelial response, and device apposition were not simulated. No angiographic or clinical follow-up data were available. The stent’s favorable computational profile therefore remains unvalidated against occlusion outcomes in this patient. The patient-specific component of the framework is likewise a single reconstruction. It demonstrates pipeline applicability to routinely available Bangladeshi clinical imaging. However, one case cannot support population-level claims about South Asian aneurysm morphology or hemodynamics. A larger patient-specific cohort is required to support such claims. These results establish a relative, hypothesis-generating comparison of device classes under matched idealized and experimental conditions. The framework extends the hemodynamic comparison to patient-specific, compliant-wall geometries and the structural analysis to cyclic fatigue loading.

## 4. Conclusion

We integrate patient-specific and idealized computational fluid dynamics (CFD), structural finite element analysis (FEA), and experimental validation using a universal testing machine (UTM) to evaluate three principal endovascular flow-diversion strategies for cerebral aneurysm treatment. We also analyze porosity and material effects.

Patient-specific CFD confirmed that the reconstructed aneurysm occupies a high-risk hemodynamic regime. Idealized device comparison demonstrated that stent-assisted diversion produces the most favorable overall hemodynamic profile among the three modeled strategies. Stent-assisted diversion was the only strategy to reduce time-averaged wall shear stress (TAWSS) relative to the untreated baseline (-21.0%). It produced the largest reductions in peak systolic velocity, wall shear stress (WSS), and vorticity at peak aneurysm-wall loading. The WEB device comes out as a strong second-line option. The coil model showed the least favorable sustained hemodynamic profile of the three, increasing peak systolic velocity and WSS relative to the untreated state rather than reducing either.

Structural FEA reproduced the clinical stiffness hierarchy we’d expect across the three candidate materials (316L stainless steel, Nitinol, and PLLA), with radial strength and stiffness decreasing consistently as porosity increased for all three. Peak equivalent stress and strain followed a non-monotonic trend, reaching minima at 55% porosity. Local strut-junction geometry, rather than porosity fraction alone, governs peak structural demand. Experimental UTM compression of FDM-printed PLA and TPU analogs confirmed this porosity ranking. The 50% porosity design showed the greatest force retention and the most delayed densification onset for both surrogate materials. PLA produced a stiffer, more brittle response; TPU produced a more gradual, elastically recoverable response. Dimensional analysis established that the scaled UTM specimens are mechanically equivalent to the native-scale devices within approximately 2.3% of normalized stiffness. A closed-form elastic-tube analysis showed that the radial compliance of all nine device configurations is negligible relative to that of the native vessel wall. This result provides an analytical basis for the rigid-wall hemodynamic assumption for the deployed devices.

These results demonstrate that a 50% porosity 316L stainless steel stent offers the greatest resistance to radial compression. A 50% porosity Nitinol stent offers the highest load-bearing capacity with more compliant, linear elastic recovery. Material selection therefore depends on the required balance between radial rigidity and elastic flexibility.

This integrated CFD-FEA-UTM framework is the first to assess hemodynamic performance across the three flow-diversion device classes. The study also characterizes representative stent prototypes through structural and mechanical testing. It is the first such framework based on South Asian patient data. The framework deploys FDM-printed analogues rather than clinical-grade implants. This design enables reproducible experiments in resource-limited research settings. Future studies must extend the hemodynamic comparison to patient-specific, compliant-wall geometries. They must also expand structural analysis to cyclic fatigue loading to strengthen translational relevance.

