## Supplementary Information for "Hemodynamic-Structural Multi-Fidelity Assessment of Endovascular Flow Diversion Strategies for Cerebral Aneurysm Treatment: An Integrated Computational and Experimental Framework"

**SI.1	Mesh Generation and Mesh Independence Study for CFD**

We discretized the computational domain with unstructured tetrahedral meshes in ANSYS. We applied 5 inflation layers along the vessel wall to resolve near-wall flow gradients. The layers used a first-layer thickness of 0.12 mm and a growth rate of 1.2. We assessed the mesh quality using the orthogonal quality and skewness metrics to ensure numerical stability. Table 1 presents a mesh independence study. We selected the mesh resolution that balanced computational accuracy and cost. We generated 11 mesh densities by progressively refining the global element size from 0.50 mm to 0.08 mm.

**Table S1.** Mesh independence study: area-weighted average wall shear stress (WSS) across mesh densities.

| Mesh Quality | Mesh Size (mm) | Element Count | Wall Shear Stress (WSS) (Pa) | Relative WSS Change (%) |
| --- | --- | --- | --- | --- |
| Coarsest | 0.50 | 66,291 | 5.91635 | 12.34 |
| Very Coarse | 0.45 | 86,614 | 5.86432 | 11.35 |
| Coarse | 0.40 | 108,832 | 5.85208 | 11.12 |
| Moderately Coarse | 0.35 | 171,410 | 5.82691 | 10.64 |
| Medium | 0.30 | 262,963 | 5.78149 | 9.78 |
| Moderately Fine | 0.25 | 441,977 | 5.73119 | 8.82 |
| Fine | 0.20 | 836,654 | 5.57538 | 5.86 |
| Slightly Finer | 0.15 | 1,928,492 | 5.36909 | 1.95 |
| Finer | 0.12 | 3,705,921 | 5.30021 | 0.64 |
| Very Fine | 0.10 | 6,339,417 | 5.26946 | 0.05 |
| Finest | 0.08 | 12,257,181 | 5.26631 | 0.00 |

We conducted transient computational fluid dynamics (CFD) simulations for each mesh under identical boundary conditions. We selected the area-weighted average wall shear stress (WSS) on the aneurysm wall as the convergence parameter. This metric quantifies mesh-refinement sensitivity and clinical relevance in aneurysm hemodynamics.

The predicted WSS decreased with increasing mesh density and approached an asymptotic value beyond approximately 3.7 million elements, as shown in Figure 1a. The relative change in WSS between the meshes containing 3,705,921 and 6,339,417 elements was 0.64%, demonstrating numerical convergence. Further refinement to 12,257,181 elements minimally changed WSS while increasing computational cost. Therefore, we selected the mesh containing 3,705,921 elements (global element size 0.12 mm) for all subsequent simulations. This mesh maintained numerical accuracy at reduced computational cost.

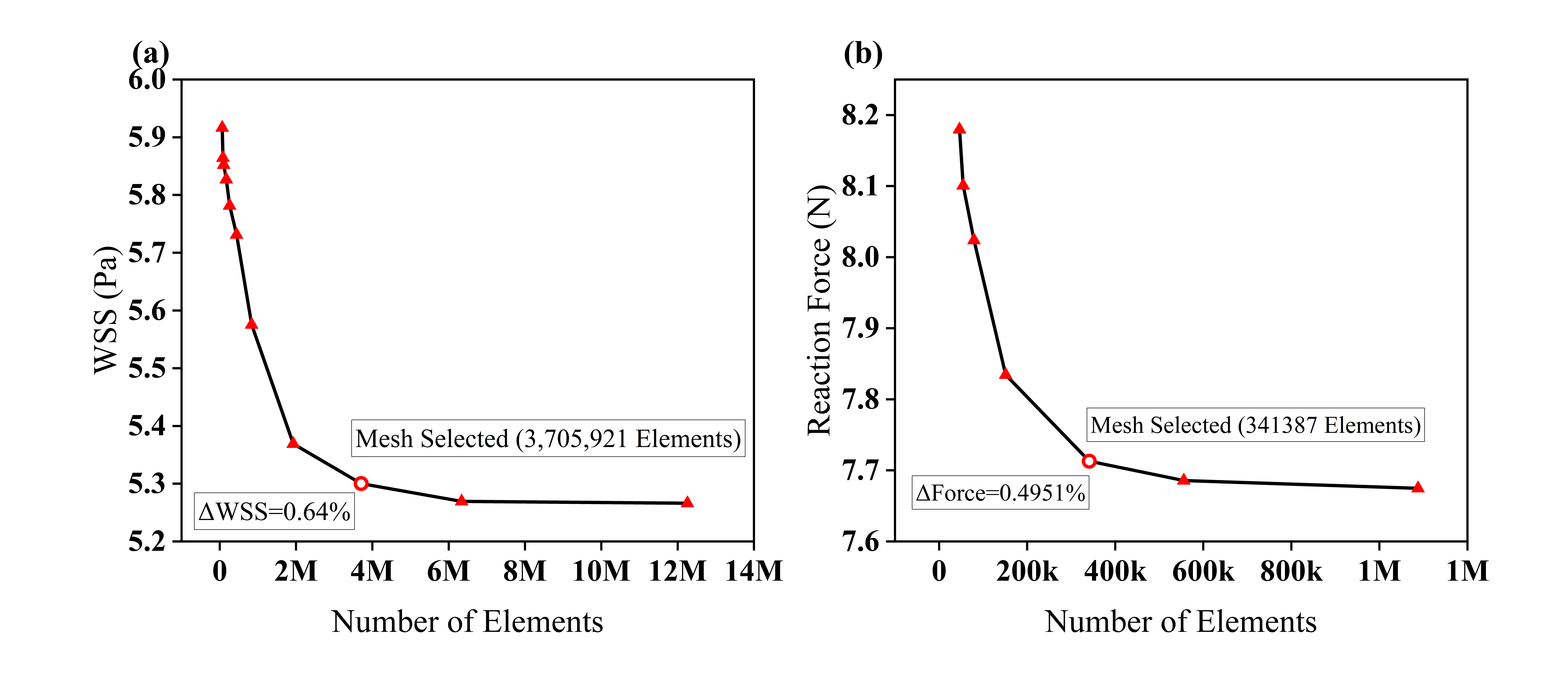
**Figure S1.** Mesh independence study showing (a) the variation of WSS with the number of computational elements and (b) reaction force as a function of the number of elements.

**SI 2.	Material Selection**

**Table S2.** Complete material parameter sets used for the three geometries in the finite element analysis; adapted from [1–3]

| Property | 316L Stainless Steel | PLLA | Nitinol |
| --- | --- | --- | --- |
| Density | 7990 kg/m^3^ | 1250 kg/m^3^ | 6450 kg/m^3^ |
| Young’s Modulus | 193,000 MPa | 3000 MPa | 50,000 MPa |
| Poisson’s Ratio | 0.31 | 0.35 | 0.33 |
| Bulk Modulus | 1.69 × 10^11^ Pa | 3.33 × 10^9^ Pa | 4.902 × 10^10^ Pa |
| Shear Modulus | 7.69 × 10^10^ Pa | 1.11 × 10^9^ Pa | 1.8797 × 10^10^ Pa |
| Tensile Yield Strength | 205 MPa | 60 MPa | 400 MPa |
| Tensile Ultimate Strength | 515 MPa | 70 MPa | N/A |

**SI 3.**	**Mesh Generation and Mesh Independence Study for FEA**

In the mesh-independence study, we evaluate seven mesh densities and compare the reaction force at maximum compression (Table 4). The results demonstrate a force difference below 0.5% relative to the finest mesh at an element size of 0.06 mm. We select this mesh (341,387 elements) for all subsequent simulations.

**Table S3.** Mesh convergence study: reaction force and percentage change relative to the finest mesh across seven element sizes.

| Mesh Quality | Element Size (mm) | Number of Elements | Reaction Force (N) | Force Change (%) |
| --- | --- | --- | --- | --- |
| Very Coarse | 0.14 | 46,370 | 8.1798 | 6.5786 |
| Coarse | 0.12 | 54,896 | 8.1008 | 5.5493 |
| Medium | 0.10 | 79,350 | 8.0238 | 4.5459 |
| Moderately Fine | 0.08 | 151,503 | 7.8345 | 2.0795 |
| Fine (selected) | 0.06 | 341,387 | 7.7129 | 0.4951 |
| Finer | 0.05 | 555,906 | 7.6857 | 0.1407 |
| Finest | 0.04 | 1,087,923 | 7.6749 | 0.0000 |

Figure S1b shows convergence. The reaction force converged with increasing element count. Over 95% of mesh elements had skewness < 0.75 and orthogonal quality > 0.1. We excluded the Very Coarse and Coarse meshes because they deviated substantially from the finest mesh. We determined convergence from the diminishing change across the medium, fine, and finest meshes. The fine mesh showed a deviation less than 0.5%.

**SI 4.	Surrogate Material Selection**

**Table S4.** Comparison between Nitinol and thermoplastic polyurethane adapted from [4,5]

| Property | Nitinol (Ni–Ti alloy) | TPU (Thermoplastic Polyurethane) | Similarity |
| --- | --- | --- | --- |
| Characteristic | Superelastic, shape-memory | Highly flexible, elastic recovery | Both exhibit elastic deformation |
| Elastic Recovery | Up to 8% recoverable strain | High elastic recovery (rubbery) | Comparable recovery response |
| Young's Modulus | 28–83 GPa (phase-dependent) | 0.002–0.05 GPa | Modulus differs substantially |
| Flexibility | Extremely compliant under load | Extremely compliant, soft | Comparable compliance |
| Deformation Behavior | High deformation with full recovery | High deformation with full recovery | Comparable deformation response |
| Fused Deposition Modeling (FDM) Printability | Not printable (metal alloy) | Excellent FDM printability with flexible filament | Printability differs |
| Biocompatibility | Clinically validated | Generally biocompatible, not implant-grade | Biocompatibility differs |
| Corrosion Resistance | Excellent (TiO₂ passive layer) | Not applicable (polymer) | Not applicable to TPU |

**Table S5.** Comparison between PLLA and PLA adapted from [6,7]

| Property | PLLA (Poly-L-lactic acid) | PLA (Polylactic acid) | Similarity |
| --- | --- | --- | --- |
| Chemical Family | Polylactic acid (L-isomer) | Polylactic acid | Same family |
| Young's Modulus | 2.7 – 4.1 GPa | 2.7 – 3.6 GPa | Comparable |
| Tensile Strength | 50 – 70 MPa | 50 – 65 MPa | Comparable |
| Density | 1.24 – 1.30 g/cm³ | 1.21 – 1.25 g/cm³ | Comparable |
| Mechanical Compliance | Semi-rigid, moderate compliance | Semi-rigid, moderate compliance | Identical |
| Biodegradability | Fully biodegradable | Slowly biodegradable | Partial similarity |
| Printability (Fused Deposition Modeling, FDM) | Difficult (hygroscopic, brittle) | Excellent, widely available | Differs |
| Biocompatibility | U.S. Food and Drug Administration (FDA)-approved, bioresorbable | Generally biocompatible | Moderate similarity |

The qualitative correspondence motivates surrogate selection. It does not justify extending porosity or material rankings from finite element analysis (FEA) to printed specimens. Sections 2.4 and 3.8 establish and verify the quantitative conditions using Buckingham Pi dimensional analysis. These conditions define mechanical equivalence between scaled PLA/TPU specimens and native-scale devices.

**SI 5.	Dimensional Analysis and Justification of Geometric Scaling**

We could not print the fused deposition modeling (FDM) specimens used for universal testing machine (UTM) validation (Section 2.3) at the 4-mm diameter of the FEA models. We therefore performed dimensional analysis to establish mechanical equivalence conditions for the scaled specimens and native-scale stent geometries. This analysis establishes the formal basis for extending conclusions from the scaled experimental tests to the clinically relevant device scale.

For the quasi-static, isothermal, parallel-plate radial compression problem, inertial and time-dependent effects are negligible. UTM uses a crosshead speed of 1 mm/min; FEA uses a single monotonic load step. Therefore, mass and time do not enter as independent dimensions; they enter only through stress. Stent response under radial compression is described by two fundamental dimensions: length [L] and stress [σ]. Table 7 lists the governing variables.

**Table S6.** Different variables with their symbol and dimensions.

| **Variable** | **Symbol** | **Dimension** |
| --- | --- | --- |
| Reaction force | *F* | [stress·length²] |
| Prescribed displacement | *δ* | [length] |
| Stent diameter | *D* | [length] |
| Stent length | *L* | [length] |
| Strut thickness | *t* | [length] |
| Young's modulus | *E* | [stress] |
| Yield strength | *σ_y_* | [stress] |
| Poisson's ratio | *ν* | [-] |
| Porosity | *φ* | [-] |

The normalized force response showing in equation (1).

$$\begin{aligned} \frac{F}{{E.D}^{2}}=\Phi\left( \frac{\delta}{D},\frac{t}{D},\frac{L}{D},\varphi,\nu,\frac{\sigma_{y}}{E} \right)\#\left( 1 \right) \end{aligned}$$

We differentiate the linear-elastic response with respect to displacement to obtain the radial-stiffness scaling law, $k=\frac{dF}{d\delta}$in equation (2).

$$\begin{aligned} k \propto E\cdot D \cdot f\left( \frac{t}{D},\frac{L}{D},\varphi,\nu,\frac{\sigma_{y}}{E} \right)\#\left( 2 \right) \end{aligned}$$

The physical content of this result is the basis for the scaling defense used: for two stent structures of arbitrary absolute size, if the normalized geometric groups $(\frac{t}{D},\frac{L}{D},\varphi)$ and the normalised material groups $(\nu,\frac{\sigma_{y}}{E})$ are matched, the normalized stiffness Π_1_ and normalized force response Φ(·) are identical regardless of the absolute diameter D. Size is not an independent variable in the governing physics of this problem; only the normalized shape of the lattice and the normalized material properties are. Table 8 below summarizes the normalized geometric groups Π_3_ and Π_4_ for the FEA models and the UTM specimens.

**Table S7.** Diameter, Length, Strut Thickness, and two dimensionless geometric variables of FEA and UTM configurations.

| Configuration | D (mm) | L (mm) | t (mm) | Π_3_=t/D | Π_4_=L/D |
| --- | --- | --- | --- | --- | --- |
| FEA model (all porosities) | 4 | 7.68 | 0.18 | 1.92 | 0.045 |
| UTM specimen (all porosities) | 25 | 48 | 1.0 | 1.92 | 0.040 |
| Geometric-similarity target for UTM | 25 | 48 | 1.125 | 1.92 | 0.045 |

**SI 6.	Coupling Flow Rate to Radial Compliance Across the Three Stent Geometries**

The CFD models in Section 2.1 and the structural analyses in Sections 2.2–2.4 examine the same three device geometries but remain mechanically uncoupled. The lumen is treated as rigid in every hemodynamic simulation. Therefore, the reported flow metrics exclude radial-compliance effects on flow rate over the cardiac cycle. This section develops a closed-form, first-order coupling between radial compliance and flow rate. The coupling applies classical flexible-tube theory. We evaluate all nine finite element analysis (FEA) configurations: 316L stainless steel, Nitinol, and PLLA at 50%, 55%, and 60% porosity. We retain the rigid-wall CFD result. We quantify the neglected compliance contribution for the three device geometries and assess differences before a full fluid-structure interaction (FSI) study.

### **SI 6.1. Elastic-Tube Theory: From Wall Modulus to Flow-Rate Sensitivity**

For steady laminar flow in a rigid tube of radius *R* driven by an axial pressure gradient, the Hagen–Poiseuille relation gives equation (3).

$$\begin{aligned} Q=\frac{{\pi R^{4}\Delta P}_{\mathrm{axial}}}{8\mu L}\#\left( 3 \right) \end{aligned}$$

the *R^4^* sensitivity that underlies the rigid-wall assumption already used in Section 2.1. A real vessel segment is not perfectly rigid. Its radius responds to the transmural (luminal) pressure. For a thin-walled elastic tube of unstressed radius *R_0_* and wall thickness *h*, Laplace’s law (hoop stress *σθ = PR/h*) combined with Hooke’s law gives the small-strain radial response in equation (4).

$$\begin{aligned} \frac{\Delta R}{R_{0}}=\frac{\Delta P.R_{0}}{E.h}\#\left( 4 \right) \end{aligned}$$

where Δ*P* is taken as the pulse-pressure amplitude already used as the outlet boundary condition in Section 2.1.2 (≈ 1.73×10^4^ Pa systolic, 1.0×10^4^ Pa diastolic ≈ 7.3×10^3^ Pa, Figure 2b) and *E* is the effective circumferential (membrane) modulus of the structure resisting distension. This is the same small-strain relation presented in equation (5) that underlies the classical Moens-Korteweg wave speed for pulse propagation in an elastic tube,

$$\begin{aligned} c_{0}=\sqrt{\frac{E.h}{{2\rho R}_{0}}}\#\left( 5 \right) \end{aligned}$$

included here because it is the standard textbook reference point for flow through flexible tubes [8,9] and because it gives a second, independently interpretable output, an equivalent pulse-wave speed, alongside the flow-rate estimate below.

Because *Q* ∝ *R^4^* at a given axial driving condition, a small fractional radius change propagates to flow rate by following equation (6).

$$\begin{aligned} \frac{\Delta Q}{Q_{0}}\approx4\left( \frac{\Delta R}{R_{0}} \right)=\frac{4\Delta P{.R}_{0}}{E.h}\#\left( 6 \right) \end{aligned}$$

that is a quasi-steady approximation. It captures the sensitivity of flow rate to a change in lumen radius but does not resolve the phase lag between the pressure and flow waveforms that a full unsteady solution would carry. The relevant dimensionless group for that phase lag is the Womersley number showing in equation (7).

$$\begin{aligned} \alpha=R_{0}\sqrt{\left( \frac{\omega\rho}{\mu} \right)}\#\left( 7 \right) \end{aligned}$$

Given the manuscript’s cardiac-cycle period (*T* = 0.8 s, *ω* = 2π/*T*) and blood density (*ρ* = 1060 kg/m³), the calculation yields *α* ≈ 0.8 with the zero-shear Carreau-Yasuda viscosity (*μ_0_* = 0.056 Pa·s) and *α* ≈ 3.1 with the high-shear-rate limit (*μ_∞_* = 0.0035 Pa·s). Both values are O(1), demonstrating that neither the fully quasi-steady (*α* ≪ 1) nor the fully inertia-dominated (*α* ≫ 1) limit strictly applies. We therefore use the flow-rate relation above as a comparative, order-of-magnitude estimate across the nine configurations, not as a phase-resolved prediction.

### **SI 6.2. Extending the Framework to the Porous Strut Lattice**

Section 2.4 from the main manuscript establishes the Buckingham Π result: the measured diametral (parallel-plate) radial stiffness of each FEA configuration scales as $k\propto E\cdot D\cdot f(\frac{t}{D},\frac{L}{D},\phi,\nu,\frac{\sigma_{y}}{E})$. The dimensionless group Π_1_ = *k/(E·D)* was evaluated for all nine combinations (Table 16). Nitinol and PLLA are modeled as linear elastic over the identical 0–1.6 mm displacement range. Their Π_1_ values demonstrate quantitative agreement at every porosity: 7.36×10^−5^ versus 7.33×10^−5^ at 50% porosity and 4.84×10^−5^ versus 4.83×10^−5^ at 60%. Therefore, *k* scales linearly with *E* at fixed geometry, as the Π-group analysis predicts. 316L SS deviates from this trend because its stiffness is extracted from a much narrower 0–0.1 mm pre-yield window (Section 2.4) and cannot be compared directly with these values.

The parallel-plate stiffness characterizes a bending-dominated mode generated by two opposing point loads. This mode represents delivery and crimping loads. Pulsatile luminal pressure loads the strut ring differently. Pressure applies a uniform circumferential load. A closed, axisymmetric ring therefore exhibits a membrane-dominated circumferential-stretching response. Classical cellular-solids theory assigns distinct stiffness–porosity scaling to these modes. Bending-dominated cellular response scales as *E_eff_* ∝ (1−φ)^2^, while stretching-dominated response scales as $E_{eff}\propto(1-\phi)$ [10]. Pressure drives this coupling. We therefore use the following effective circumferential modulus at equation (8).

$$\begin{aligned} E_{eff}{=E}_{solid}.\left( 1-\varphi\right)\#\left( 8 \right) \end{aligned}$$

We evaluate this modulus at the as-built porosities reported in Section 2.2.1 (50.2%, 55.9%, and 59.3%; nominal targets: 50%, 55%, and 60%). We combine it with the design strut thickness (*h* = 0.18 mm) and the stent radius (*R_0_* = 2 mm, matching the 4 mm parent-vessel diameter in Table 2). The pressure loading produces minimal strains and does not engage Nitinol’s superelastic transformation (Table 9). Therefore, the linear modulus tabulated in Table 3 is physically appropriate. The full-scale FEA compression analysis required the nonlinear constitutive model.

***SI 6.3.	Applied Results Across Nine Configurations***

Table 8 reports the equations above for all nine material-porosity combinations. It also includes three reference rows for the native (untreated) arterial/aneurysmal wall. These rows use representative literature values (*E* ≈ 0.4–2 MPa [11], report intracranial arteries as significantly stiffer than extracranial arteries of the same subject, consistent with values toward the upper end of this range being appropriate for a cerebral vessel). Each reference case uses a wall thickness equal to 10% of the parent-vessel diameter. This parameterization is standard in image-based cerebrovascular FSI modeling when patient-specific wall thickness is unavailable.

**Table S8.** Predicted radial distension, flow-rate modulation and equivalent pulse-wave speed for the nine FEA material-porosity configurations and three native-wall reference cases.

| Configuration | Porosity | *E*_eff_ (MPa) | *ΔR*/*R*_0_ (%) | *ΔQ*/*Q*_0_ (%) | *c*_0_ (m/s) |
| --- | --- | --- | --- | --- | --- |
| 316L SS | 50.2% | 96,114 | 8.44 × 10^−5^ | 3.38 × 10^−4^ | 2020 |
|  | 55.9% | 85,113 | 9.53 × 10^−5^ | 3.81 × 10^−4^ | 1901 |
|  | 59.3% | 78,551 | 1.03 × 10^−4^ | 4.13 × 10^−4^ | 1826 |
| Nitinol | 50.2% | 24,900 | 3.26 × 10^−4^ | 1.30 × 10^−3^ | 1028 |
|  | 55.9% | 22,050 | 3.68 × 10^−4^ | 1.47 × 10^−3^ | 968 |
|  | 59.3% | 20,350 | 3.99 × 10^−4^ | 1.59 × 10^−3^ | 929 |
| PLLA | 50.2% | 1,494 | 5.43 × 10^−3^ | 2.17 × 10^−2^ | 252 |
|  | 55.9% | 1,323 | 6.13 × 10^−3^ | 2.45 × 10^−2^ | 237 |
|  | 59.3% | 1,221 | 6.64 × 10^−3^ | 2.66 × 10^−2^ | 228 |
| Native wall (low, 0.4 MPa) | N/A | 0.4 | 9.13 | 41.8 | 6.1 |
| Native wall (mid, 1.0 MPa) | N/A | 1.0 | 3.65 | 15.4 | 9.7 |
| Native wall (high, 2.0 MPa) | N/A | 2.0 | 1.82 | 7.5 | 13.7 |

Across all nine device configurations, the predicted radius change never exceeds 7×10^−3^%. This maximum occurs for PLLA at 59.3% porosity, its most compliant configuration. The minimum is 8×10^−5^% for 316L SS at 50% porosity. Even the 64-fold spread in solid-material Young’s modulus between 316L SS and PLLA collapses to less than a two-order-of-magnitude spread in Δ*R*/*R_0_* once combined with the porosity and geometry terms. Both extremes remain three to five orders of magnitude below any distension that would be resolvable in the CFD mesh used in Section 2.1 from the manuscript. These results demonstrate that the deployed stent can be treated as rigid in the hemodynamic simulations across the applied pulse-pressure range. This conclusion is independent of the three candidate materials. None of the nine device configurations behaves as a mechanically flexible tube.

**SI 7.	Hemodynamic Parameters**

**Table S9.** Instantaneous hemodynamic parameters at peak systole (t = 0.07 s).

| Parameter | Untreated | Stent | Coil | WEB |
| --- | --- | --- | --- | --- |
| Velocity (m/s) | 0.2565 | 0.2141 | 0.2964 | 0.2122 |
| Pressure (Pa) | 11700.5 | 11771.0 | 11664.3 | 11726.1 |
| WSS (Pa) | 10.060 | 8.569 | 10.674 | 10.056 |
| Vorticity | 2731.16 | 2317.60 | 2896.55 | 2728.90 |

**Table S10.** Instantaneous hemodynamic parameters at end diastole (t = 0.78 s).

| Parameter | Untreated | Stent | Coil | WEB |
| --- | --- | --- | --- | --- |
| Velocity (m/s) | 0.0181 | 0.01871 | 0.01765 | 0.01713 |
| Pressure (Pa) | 12700.2 | 12699.3 | 12700.9 | 12697.8 |
| WSS (Pa) | 0.165 | 0.146 | 0.1373 | 0.1867 |
| Vorticity | 26.682 | 23.553 | 21.668 | 25.921 |

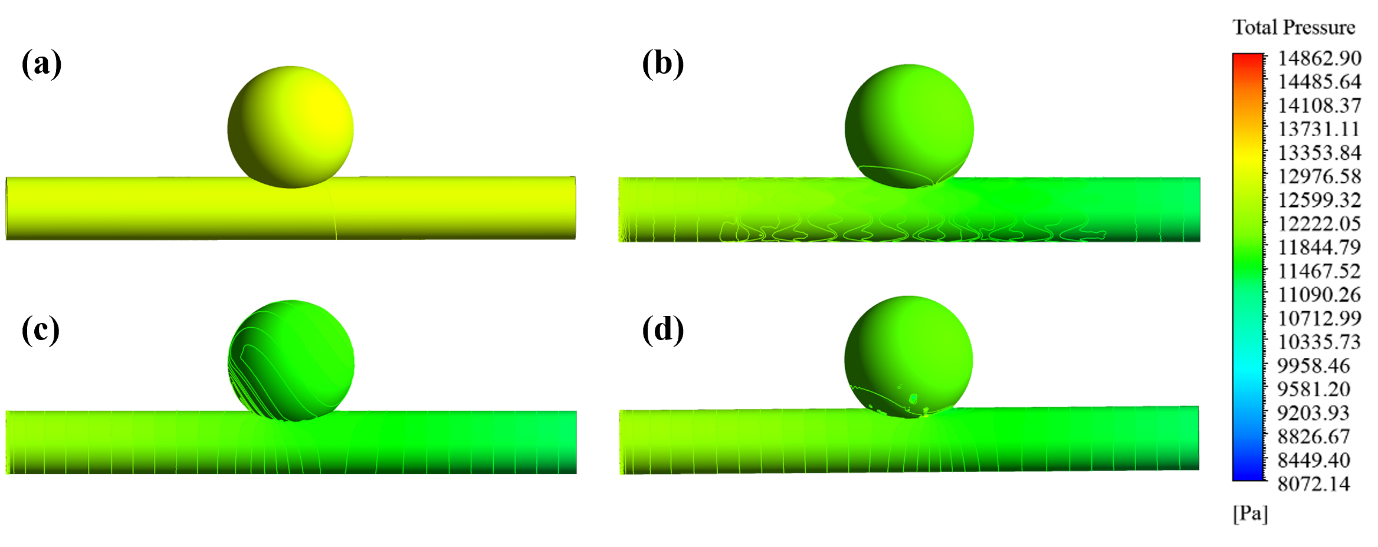
**Figure S2.** Peak-systolic total pressure contours for (a) untreated, (b) stent, (c) coil, and (d) WEB models.

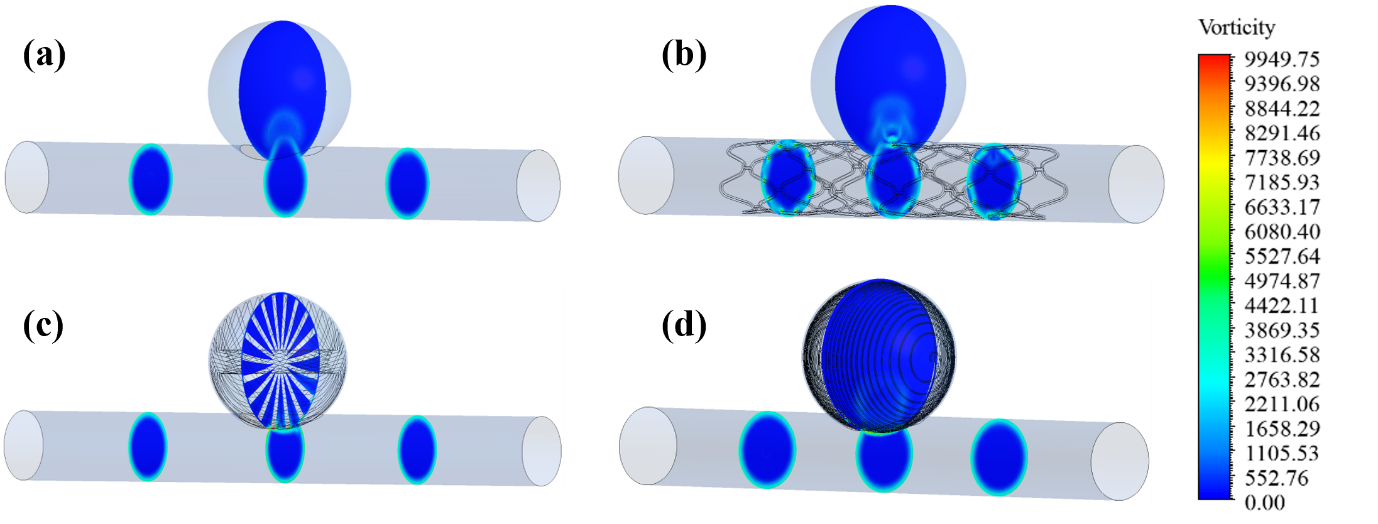

**Figure S3.** Peak-systolic vorticity contours on parent-vessel cross-sections for (a) untreated, (b) stent, (c) coil, and (d) WEB models.

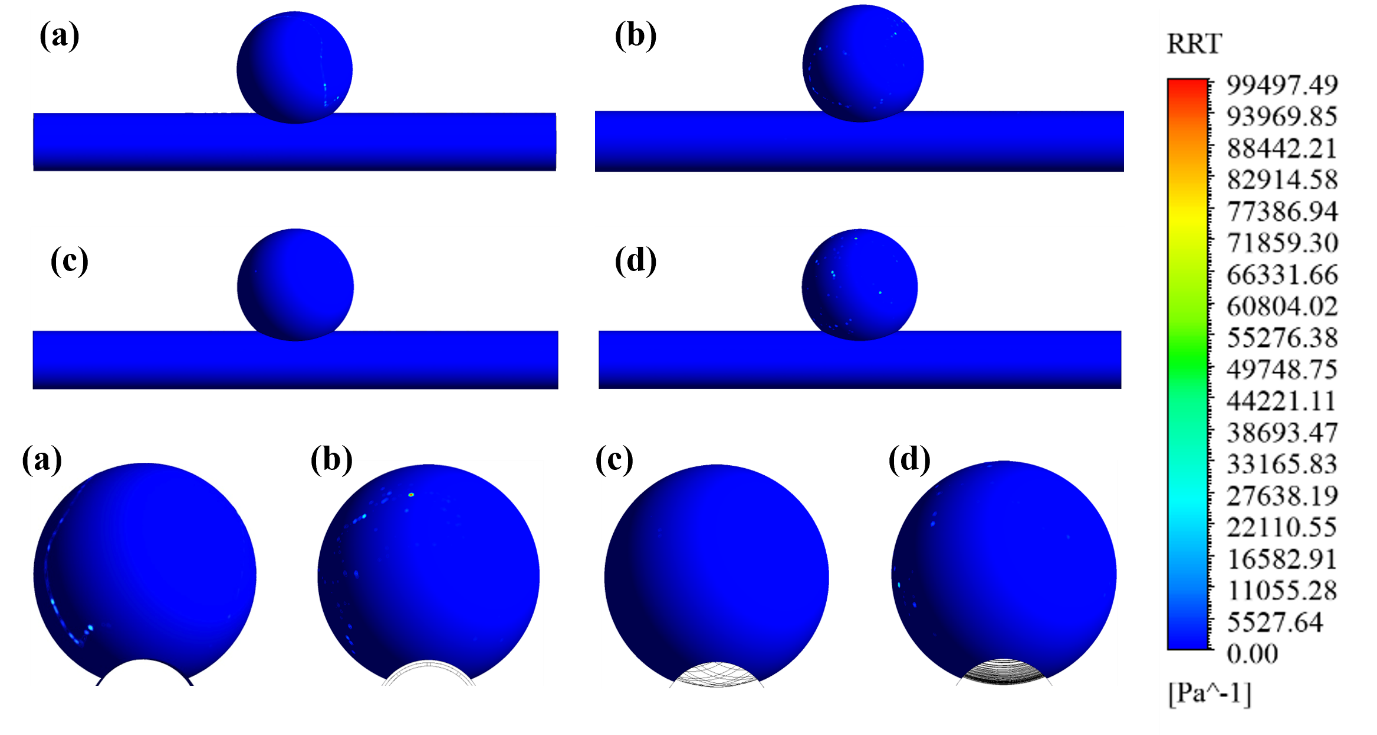

**Figure S4.** Relative residence time (RRT) distribution for (a) untreated, (b) stent, (c) coil, and (d) WEB models (side and dome views).

**SI 8.	Finite Element Analysis**

**Table S11.** Summary of maximum equivalent stress, maximum equivalent strain, reaction force at 1.6 mm displacement and radial stiffness for all porosity and material combinations.

| Material | Porosity (%) | Maximum Equivalent Stress (MPa) | Maximum Equivalent Strain | Reaction Force at 1.6mm (N) | Radial Stiffness (N/mm) |
| --- | --- | --- | --- | --- | --- |
| 316L Stainless Steel | 50 | 436.82 | 2.25E-03 | 7.72 | 45.22 |
|  | 55 | 391.64 | 2.03E-03 | 7.28 | 29.26 |
|  | 60 | 530.75 | 3.18E-03 | 5.15 | 16.27 |
| Nitinol | 50 | 2915.1 | 5.95E-02 | 23.59 | 14.71 |
|  | 55 | 2510.2 | 5.04E-02 | 17.69 | 10.97 |
|  | 60 | 4009.8 | 9.31E-02 | 15.55 | 9.67 |
| PLLA | 50 | 174.56 | 5.93E-02 | 1.41 | 0.88 |
|  | 55 | 150.36 | 0.05031 | 1.05 | 0.65 |
|  | 60 | 233.24 | 9.15E-02 | 0.93 | 0.58 |

**SI 9.	Dimensional Scaling Verification**

**Table S12.** Π_1_ values for different materials and porosities of both FEA and UTM settings.

| Material | Porosity (%) | k (N/mm) | E (MPa) | Π1 = k/(E·D) |
| --- | --- | --- | --- | --- |
| 316L SS | 50 | 45.22 | 193,000 | 5.86E-5 |
|  | 55 | 29.26 |  | 3.79E-5 |
|  | 60 | 16.27 |  | 2.11E-5 |
| Nitinol | 50 | 14.71 | 50,000 | 7.36E-5 |
|  | 55 | 10.97 |  | 5.49E-5 |
|  | 60 | 9.67 |  | 4.84E-5 |
| PLLA | 50 | 0.88 | 3,000 | 7.33E-5 |
|  | 55 | 0.64 |  | 5.33E-5 |
|  | 60 | 0.58 |  | 4.83E-5 |
| PLA (UTM surrogate) | 50 | 15.68 | 2800 | 22.4E-5 |
|  | 55 | 4.739 |  | 6.77E-5 |
|  | 60 | 3.458 |  | 4.94E-5 |
| TPU (UTM surrogate) | 50 | 0.29 | 50 | 23.2E-5 |
|  | 55 | 0.0985 |  | 7.88E-5 |
|  | 60 | 0.0696 |  | 5.57E-5 |
